# Structural basis for activation of Arf1 at the Golgi complex

**DOI:** 10.1101/2022.05.04.490673

**Authors:** Arnold J. Muccini, Margaret A. Gustafson, J. Christopher Fromme

**Affiliations:** Department of Molecular Biology and Genetics, Weill Institute for Cell and Molecular Biology, Cornell University, Ithaca, NY 14853 USA

## Abstract

The Golgi complex is the central sorting station of the eukaryotic secretory pathway. Traffic through the Golgi requires activation of Arf GTPases that orchestrate cargo sorting and vesicle formation by recruiting an array of effector proteins. Arf activation and Golgi membrane association is controlled by large guanine nucleotide exchange factors (GEFs) possessing multiple conserved regulatory domains. Here we present cryoEM structures of full-length Gea2, the yeast paralog of the human Arf-GEF GBF1, that reveal the organization of these regulatory domains and explain how Gea2 binds to the Golgi membrane surface. We find the GEF domain adopts two different conformations compatible with different stages of the Arf activation reaction. The structure of a Gea2-Arf1 activation intermediate suggests the movement of the GEF domain primes Arf1 for membrane insertion upon GTP binding. We propose that conformational switching of Gea2 during the nucleotide exchange reaction promotes membrane insertion of Arf1.

## Introduction

The endomembrane system provides essential compartmentalization for all eukaryotic cells. Most transmembrane and lumenal proteins are synthesized at the ER and then travel through the secretory pathway to reach their target organelle. At the center of the secretory pathway is the Golgi complex, which modifies secretory proteins and serves as a trafficking hub. Arf1 and its close paralogs are essential regulators of cargo sorting and vesicle formation at the Golgi complex that function by recruiting a large number of prominent effectors including COPI/coatomer, clathrin cargo adaptors, lipid signaling enzymes, vesicle tethers, and regulators of other pathways (Adarska et al., 2021; Cherfils, 2014; Donaldson and Jackson, 2011;Gillingham and Munro, 2007). Arf1 is a GTPase, cycling between an inactive GDP-bound state and an active GTP-bound state (Kahn and Gilman, 1986). Arf1 possesses an N-terminal myristoylated amphipathic helix that anchors it to the Golgi membrane (Haun et al., 1993; Kahn et al., 1988). When GDP-bound, this membrane-binding feature is masked and Arf1 is cytosolic. When Arf1 is activated to its GTP-bound state, a change in conformation exposes the myristoylated amphipathic helix, resulting in stable membrane-association (Amor et al., 1994;Antonny et al., 1997; Franco et al., 1995; Goldberg, 1998). The active conformation of Arf1 is therefore required to recruit its numerous effectors to the Golgi membrane surface.

Arf1 activation in cells requires nucleotide exchange by specific guanine-nucleotide exchange factors (GEFs). Arf1 is activated at the Golgi complex by at least two distinct but related Arf-GEFs, GBF1 and BIG1/2 (Claude et al., 1999; Togawa et al., 1999). The budding yeast homolog of BIG1/2 is Sec7, which localizes to late Golgi compartments and activates Arf1 to control trafficking to endosomes, lysosomes, earlier Golgi compartments, and the plasma membrane (Franzusoff et al., 1991; Novick et al., 1981). The budding yeast homologs of GBF1, named Gea1/2, localize to early and medial Golgi compartments where Arf1 activation orchestrates the formation of COPI vesicles destined for the endoplasmic reticulum and earlier Golgi compartments (Gustafson and Fromme, 2017; Peyroche et al., 1996; Spang et al., 2001).

The Golgi Arf-GEFs share a homologous catalytic GEF domain, referred to as a “Sec7” domain, with members of other Arf-GEF families (Casanova, 2007). The structural and biochemical basis for nucleotide exchange by Sec7 GEF domains is well-established and involves remodeling of the Arf1 nucleotide-binding site by interaction with the GEF (Goldberg,1998; Renault et al., 2003). The ARNO/cytohesin/Grp1 and BRAG/IQSec7 Arf-GEFs possess structurally characterized plekstrin homology (PH) domains that direct membrane binding and regulation of GEF activity (Aizel et al., 2013; Cronin et al., 2004; Das et al., 2019; DiNitto et al.,2007; Malaby et al., 2018). In contrast, the Golgi-localized “large” Arf-GEFs do not contain PH domains and instead contain multiple regulatory domains that are conserved across species but are not found in other proteins (Bui et al., 2009; Mouratou et al., 2005). Previous studies have dissected the biochemical and cell biological roles of these regulatory domains and identified which domains are required for Golgi membrane binding and activation of Arf1 (Bouvet et al.,2013; Christis and Munro, 2012; Gustafson and Fromme, 2017; Meissner et al., 2018; Pocognoni et al., 2018; Richardson and Fromme, 2012; Richardson et al., 2012). Structures are available for the N-terminal ‘DCB-HUS’ domains in isolation (Galindo et al., 2016; Richardson et al., 2016; Wang et al., 2016), but the lack of structural information for the full-length proteins has prevented an understanding of how the regulatory domains function together with the GEF domain during Arf1 activation.

Here we present cryoEM structures of full-length Gea2 and a Gea2-Arf1 activation intermediate. These structures reveal the organization of the regulatory domains within the Gea2 dimer. We identify two new conserved structural elements in Gea2: an amphipathic helix between the HDS1 and HDS2 domains that is required for membrane binding, and an ordered linker between the GEF and HDS1 domains. Unexpectedly, the GEF domain of Gea2 adopts two conformational states. Structural analysis indicates that the GEF-HDS1 linker plays a role in conformational switching: the “closed” state of the GEF domain is compatible with initial binding to Arf1-GDP but incompatible with subsequent binding to nucleotide-free Arf1 due to a steric clash between nucleotide-free Arf1 and the linker. The structural data therefore suggest that the Arf1 nucleotide exchange reaction involves conformational change of its GEF from the closed state to the “open” state. Based on the orientation of Gea2 on the membrane, this GEF conformational change appears to directly couple Arf1 activation to membrane insertion.

## Results

### Architecture of the Gea2 homodimer

Gea2 and its paralogs possess an N-terminal DCB-HUS regulatory domain and C-terminal HDS1, HDS2, and HDS3 regulatory domains (Mouratou et al., 2005; Richardson et al.,2016) (Fig. 1A). We produced full-length *S. cerevisiae* Gea2 by overexpression in *Pichia pastoris* (Fig. S1A) and determined its structure using cryoEM (Figs. S2, S3). 3D classification of the particles revealed three distinct conformations of Gea2 homodimers that differed only in the positioning of the GEF domain, with each monomer adopting either a “closed” or “open” position relative to the regulatory domains (Fig. S2). Based on the relative numbers of particle images that sorted into each of these three classes (~30% “closed/closed”, ~30% “open/open”, and ~40% “closed/open”), the conformation adopted by each monomer within the dimer appears to be largely independent of that of its binding partner. We took advantage of the 2-fold symmetry of the Gea2 homodimer by using symmetry expansion and focused refinements during data processing (see Methods) to obtain higher resolution maps for the closed and open monomers and for the three different dimeric states (Figs. 1B and S4). These maps were then used to build and refine atomic models (Figs. 1C-E and S4, Table 1). We begin our description of the structure using the “closed/open” dimer as it exhibits both the closed (Fig. 1D) and open (Fig. 1E) states of the GEF domain.

**Figure 1.**
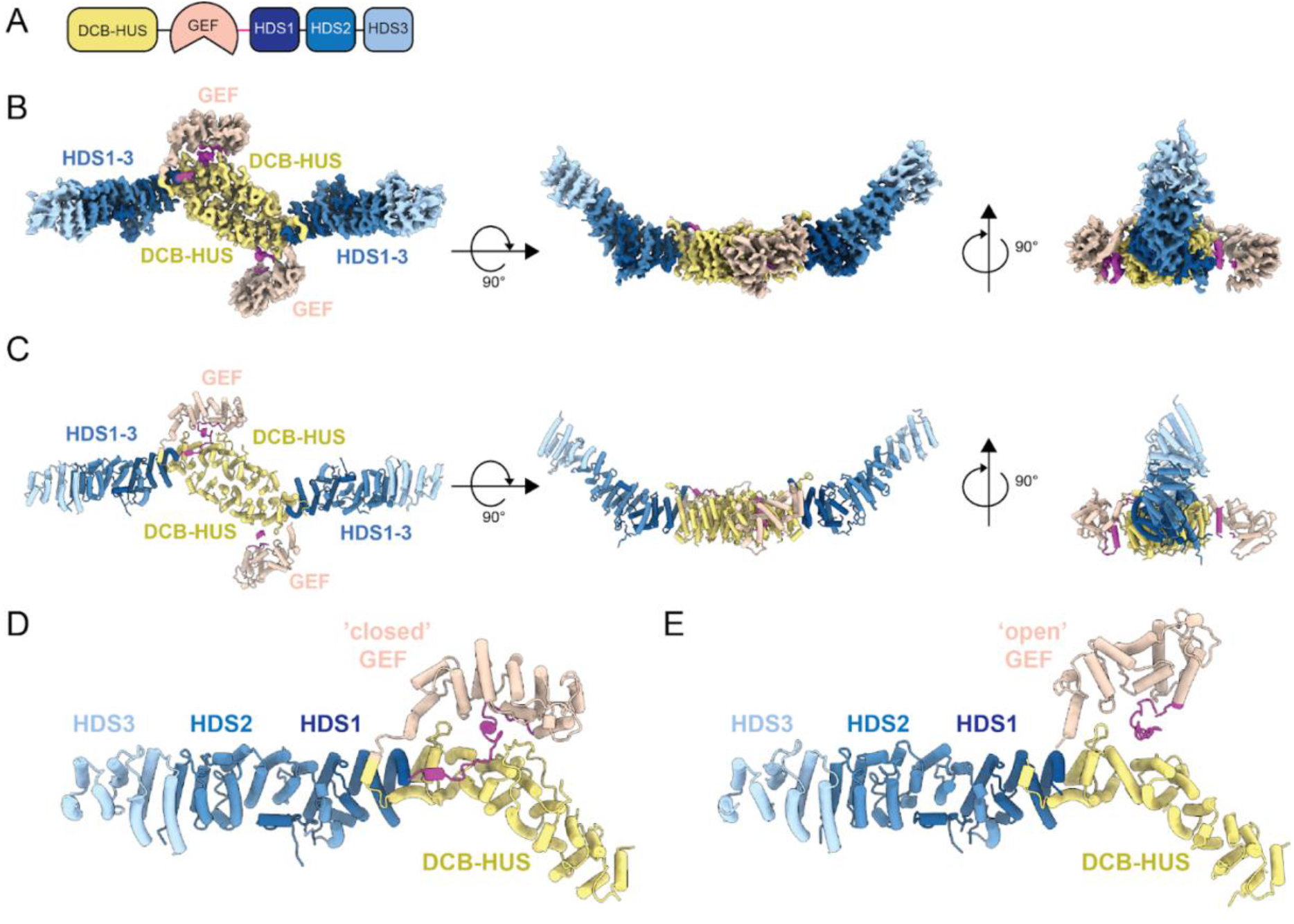
Structure of Gea2 determined by cryoEM. A) Schematic of Gea2 primary structure indicating conserved domains. DCB: dimerization and cyclophilin binding; HUS: homology upstream of Sec7; GEF: guanine nucleotide exchange factor (aka “Sec7 domain”); HDS: homology downstream of Sec7. B) CryoEM density of the Gea2 dimer in its closed/open conformation. One monomer adopts an open conformation of the GEF domain, the other monomer adopts a closed conformation. The GEF-HDS1 linker is colored magenta. C) Atomic model of the Gea2 dimer, shown in cartoon depiction. D) Close-up view of the closed monomer. E) Close-up view of the open monomer.

**Table 1.** CryoEM data collection, processing and model validation statistics.

|  | Gea2<br>closed/open<br>(composite<br>map) | Gea2<br>closed/open<br>(consensus<br>map) | Gea2<br>closed/closed<br>(composite<br>map) | Gea2<br>closed/closed<br>(consensus<br>map) | Gea2<br>open/open<br>(composite<br>map) | Gea2<br>open/open<br>(consensus<br>map) | Gea2-Arf1<br>complex<br>(composite<br>map) | Gea2-Arf1<br>complex<br>(consensus<br>map) |
| --- | --- | --- | --- | --- | --- | --- | --- | --- |
| Nominal<br>magnification | 63000 |  |  |  |  |  |  |  |
| Voltage (kV) | 200 |  |  |  |  |  |  |  |
| Total dose (e-<br>/Å<sup>2</sup>) | 50 |  |  |  |  |  |  |  |
| Defocus range<br>(µm) | -1.0 to -2.0 |  |  |  |  |  |  |  |
| Pixel Size (Å) | 1.29 |  |  |  |  |  | 1.30 |  |
| Symmetry<br>imposed | N/A | C1 | N/A | C2 | N/A | C2 | N/A | C2 |
| Particle images |  | 101014 |  | 83665 |  | 74385 |  | 391360 |
| Map resolution,<br>0.143-FSC (Å) |  | 4.7 |  | 5.1 |  | 5.1 |  | 4.23 |
| Map<br>sharpening B<br>factor | N/A | -76 | N/A | -83 | N/A | -73 | N/A | -167 |
| # atoms | 38358 | N/A | 39735 | N/A | 36590 | N/A | 42915 | N/A |
| # protein<br>residues | 2360 |  | 2448 |  | 2242 |  | 2647 |  |
| B factor,<br>protein | 186.<br>1 |  | 145.70 |  | 124.29 |  | 124.83 |  |
| R.m.s<br>deviation, bond<br>length (Å) | 0.006 |  | 0.006 |  | 0.006 |  | 0.006 |  |
| R.m.s.<br>deviation, bond<br>angles | 1.039 |  | 0.969 |  | 0.976 |  | 0.944 |  |
| MolProbity<br>score | 1.68 |  | 1.62 |  | 1.49 |  | 1.57 |  |
| Clashscore | 5.26 |  | 4.96 |  | 3.69 |  | 5.13 |  |
| Poor rotamers<br>(%) | 0.68 |  | 0.48 |  | 0.38 |  | 0.04 |  |
| Ramachandran<br>plot |  |  |  |  |  |  |  |  |
| Favored (%) | 94.03 |  | 94.85 |  | 95.28 |  | 95.68 |  |
| Allowed (%) | 5.89 |  | 5.11 |  | 4.54 |  | 4.28 |  |
| Disallowed (%) | 0.09 |  | 0.04 |  | 0.18 |  | 0.04 |  |

The HDS1, 2, and 3 domains form an extended helical repeat structure that is contiguous with the DCB-HUS domain, such that the HDS3 domains of each monomer lie at the distal ends of the homodimer (Fig. 1B,C). The GEF domain lies adjacent to the HUS domain and is connected to the HUS and HDS1 domains through ordered linker regions (Fig. S5). The “HUS box”, which is a conserved region near the C-terminal end of the HUS domain (Mouratou et al., 2005), interacts directly with the HUS-GEF linker, which is simply an extension of the first α-helix of the GEF domain (Fig. S5E-G). Temperature sensitive mutations have been identified in the region surrounding the HUS box (Park et al., 2005), lending support to the importance of this interaction. The linker that connects the GEF domain to the HDS1 domain (GEF-HDS1 linker) comprises ~45 conserved ordered residues and is discussed in detail further below (Fig. S5A-C).

Dimerization occurs through extensive hydrophobic, polar, and electrostatic interactions between the DCB-HUS domains of each monomer (Fig. 2A-F), consistent with the established role of this domain for dimerization of Gea2/GBF1 homologs (Bhatt et al., 2016; Grebe et al.,2000; Ramaen et al., 2007). The fold of the Gea2 DCB-HUS domain is quite similar to that of the distinct Arf-GEF Sec7 (Richardson et al., 2016), although this domain does not appear to mediate dimerization of Sec7. Previous studies identified substitution mutations in the DCB subdomain of GBF1 that disrupted its dimerization, in residues corresponding to K124 and D163 in Gea2 (Bhatt et al., 2016; Ramaen et al., 2007). Examination of the dimerization interface indicates that K124 is involved in favorable interactions between monomers (Fig. 2E). Therefore the observed dimerization interface is supported by these published functional results and likely conserved across Gea2/GBF1 paralogs in different species.

**Figure 2.**
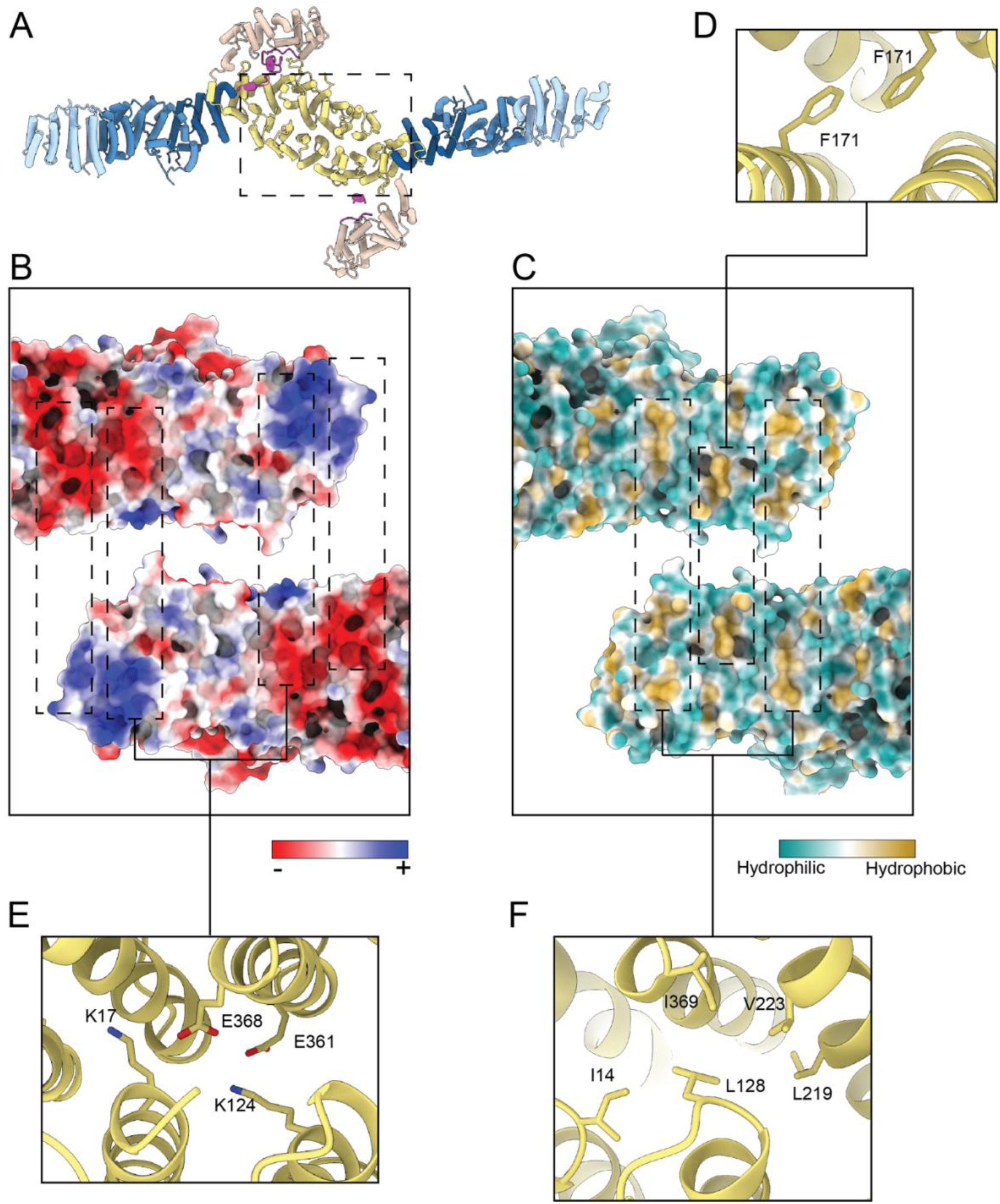
Gea2 dimerizes via the DCB-HUS domains. A) Gea2 dimer with a dashed box indicating the region depicted in (B) and (C). B) View of the dimerization interface peeled apart and colored by calculated charge potential. C) View of the dimerization interface peeled apart and colored by hydrophobicity. D-F) Close-up views highlighting residues involved in the dimerization interface.

### Gea2 binds to the Golgi via a conserved amphipathic helix

Several Arf-GEFs possess pleckstrin homology (PH) domains that bind to membranes via specific interactions with phosphoinositide lipids (Casanova, 2007). The Golgi Arf-GEFs do not possess a PH domain and although the HDS1-3 domains are known to be important for Golgi localization of Gea1/2 and GBF1 (Bouvet et al., 2013; Gustafson and Fromme, 2017;Meissner et al., 2018; Pocognoni et al., 2018), their membrane binding mechanism is unknown.

Analysis of the Gea2 cryoEM structures revealed the presence of an unstructured but conserved sequence in the linker between the HDS1 and HDS2 domains (Fig. 3A-C). This sequence is predicted to form an amphipathic helix by both secondary and tertiary sequence prediction methods (Fig. 3D). We reasoned that its conservation, position, and flexible connection to the rest of the protein made this sequence a strong candidate for a membrane-inserting amphipathic helix (Drin and Antonny, 2010). We note that this helix is distinct from amphipathic helices in the HDS1-2 domains previously proposed by other groups to be important for membrane-binding. Our structural data indicate that the amphipathic helices previously studied by others are instead part of the core helical repeat structure of these domains. As the hydrophobic faces of these helices are buried within the hydrophobic protein interior, they are unavailable for membrane interaction.

**Figure 3.**
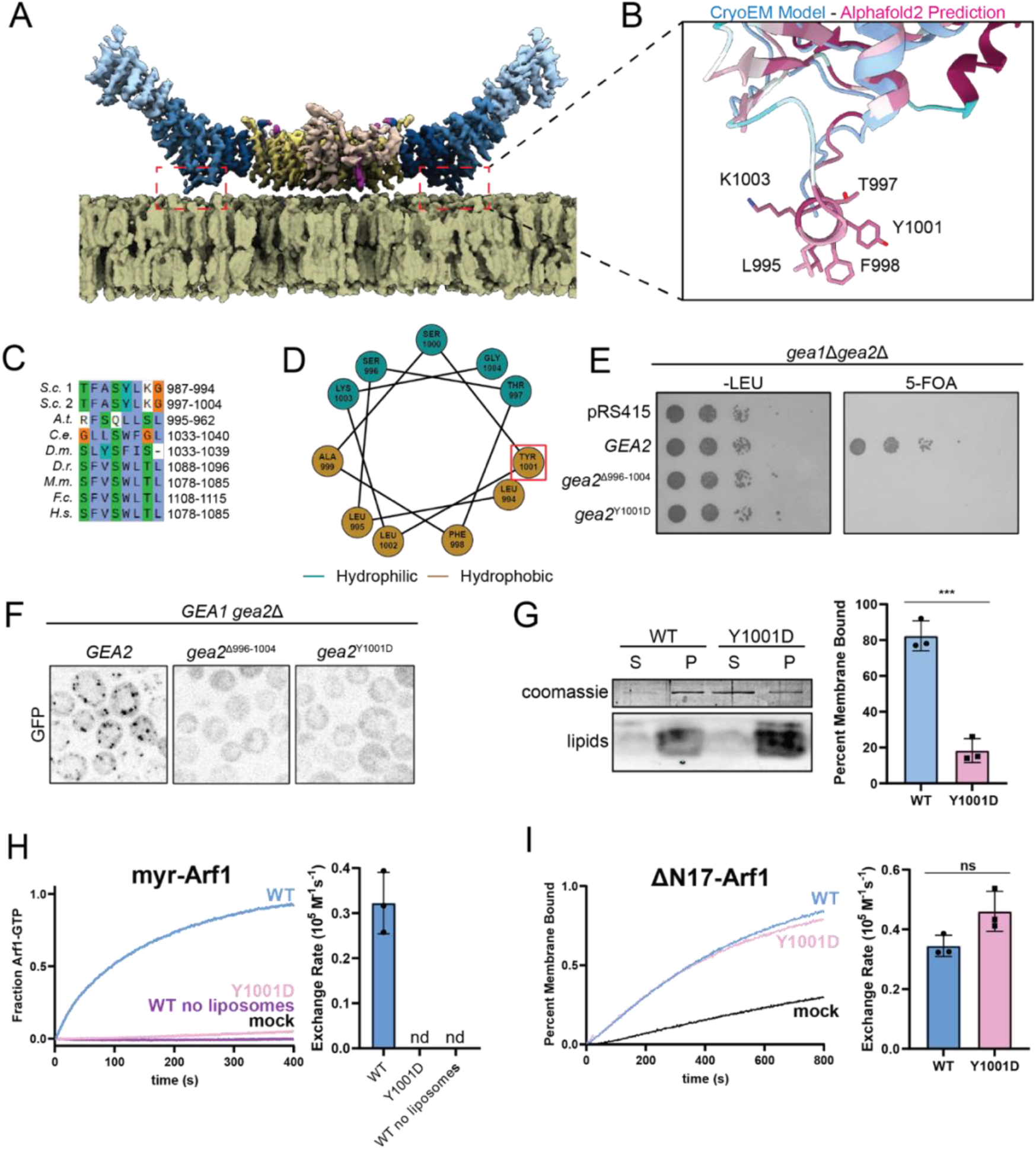
A conserved amphipathic α-helix mediates Gea2 membrane binding. A) Gea2 depicted on a modeled membrane surface. B) Close-up view of the amphipathic helix predicted by both secondary and tertiary structure prediction methods, but absent from the experimentally determined cryoEM density. The structural model determined by cryoEM is superimposed onto the AlphaFold prediction (Jumper et al., 2021). The AlphaFold prediction is colored by conservation with dark red representing the most conserved residues and cyan representing the least conserved residues. C) Sequence alignment highlighting conservation of the helix, colors highlight conserved residues based on their biochemical properties. D) Helical wheel indicating the amphipathic nature of the helix. Red box indicates Tyr residue mutated for functional experiments. E) *GEA2* complementation test (plasmid shuffling). F) Localization analysis of Gea2 and amphipathic helix mutants. G) *In vitro* membrane-binding assay (liposome pelleting) using purified proteins and synthetic liposomes. S, supernatant; P, pellet. H) In vitro GEF activity assay using purified proteins, the myristoylated-Arf1 substrate, and synthetic liposomes. nd = not detectable. I) In vitro GEF activity assay using purified proteins and the ΔN17-Arf1 substrate without liposomes.

To test the role and importance of this newly identified amphipathic α-helix, we produced two different mutants of Gea2, one in which this helix was removed (Δ996-1004) and another in which a conserved Tyr residue was substituted with Asp (Y1001D). We found that both of these mutants lost their ability to support cell growth, despite being expressed at endogenous levels (Figs. 3E, S1B). We also observed that these mutant proteins lost their localization to the Golgi complex, localizing instead to the cytoplasm (Fig. 3F). These results indicate that this conserved amphipathic helix is required for Golgi membrane association *in vivo*.

To determine whether this amphipathic helix is involved in direct interaction between Gea2 and the membrane surface, we purified the Gea2 Y1001D mutant protein (Fig. S1C) and tested its ability to interact with liposome membranes *in vitro*. Using a lipid mix that wild-type Gea2 associates with robustly, we found that the Y1001D mutant protein exhibited a dramatic reduction in membrane-binding capability *in vitro* (Fig. 3G). This indicates that the amphipathic helix is directly involved in Gea2 membrane binding.

To determine whether the amphipathic helix is required for membrane-proximal Arf1 activation, we employed an established *in vitro* GEF assay for Gea2 (Gustafson and Fromme,2017). We found that purified Gea2 Y1001D was well behaved biochemically but unable to activate full-length myristolated-Arf1 on liposome membranes (Figs. 3H, S1D). A similar lack of activation was seen when liposomes were omitted from reactions with wild-type Gea2 (Fig. 3H). Importantly, Gea2 Y1001D retained robust GEF activity towards ΔN17-Arf1 in the absence of liposome membranes (Fig. 3I, S1E). ΔN17-Arf1 is a truncated form of Arf1 that lacks its N-terminal amphipathic helix and therefore does not need to insert into membranes in order to be activated (Kahn et al., 1992; Paris et al., 1997). These results indicate that the Gea2 amphipathic helix is specifically required for activating Arf1 on the membrane surface.

Taken together our results indicate that Gea2 uses the conserved amphipathic helix in the HDS1-HDS2 linker to bind to the Golgi membrane surface in order to activate Arf1. The dimeric nature of Gea2 enables us to model its orientation on the membrane with high confidence (Fig. 3A). These findings also highlight how Arf1 activation and insertion of its myristoylated N-terminal helix into a membrane are intimately coupled.

### Gea2 adopts an open conformation when bound to nucleotide-free Arf1

To further investigate the role of the regulatory domains in modulating the action of the GEF domain, we trapped the Gea2-Arf1 nucleotide-free activation intermediate (Fig. S1F) and determined its structure by cryoEM (Figs. 4A-C and S6, Table 1). The conformation of nucleotide-free Arf1 in our full-length Gea2-Arf1 complex structure was nearly identical to that of nucleotide-free Arf1 when bound to the isolated Gea2 GEF domain determined previously by X-ray crystallography (Fig. S7A) (Goldberg, 1998). Strikingly, a closed conformation of the GEF domain was not observed in the Gea2-Arf1 complex cryoEM data; instead the position of the Arf1-bound GEF domain was similar to that of the open conformation observed in the absence of Arf1 (Figs. 4D, S6, and S7B). We note that structural predictions of Gea2, its yeast paralog Gea1, and its human homolog GBF1 each adopt the closed conformation (Fig. S7C). These structural results suggest that binding to nucleotide-free Arf1 enforces an open conformation of the Gea2 GEF domain.

**Figure 4.**
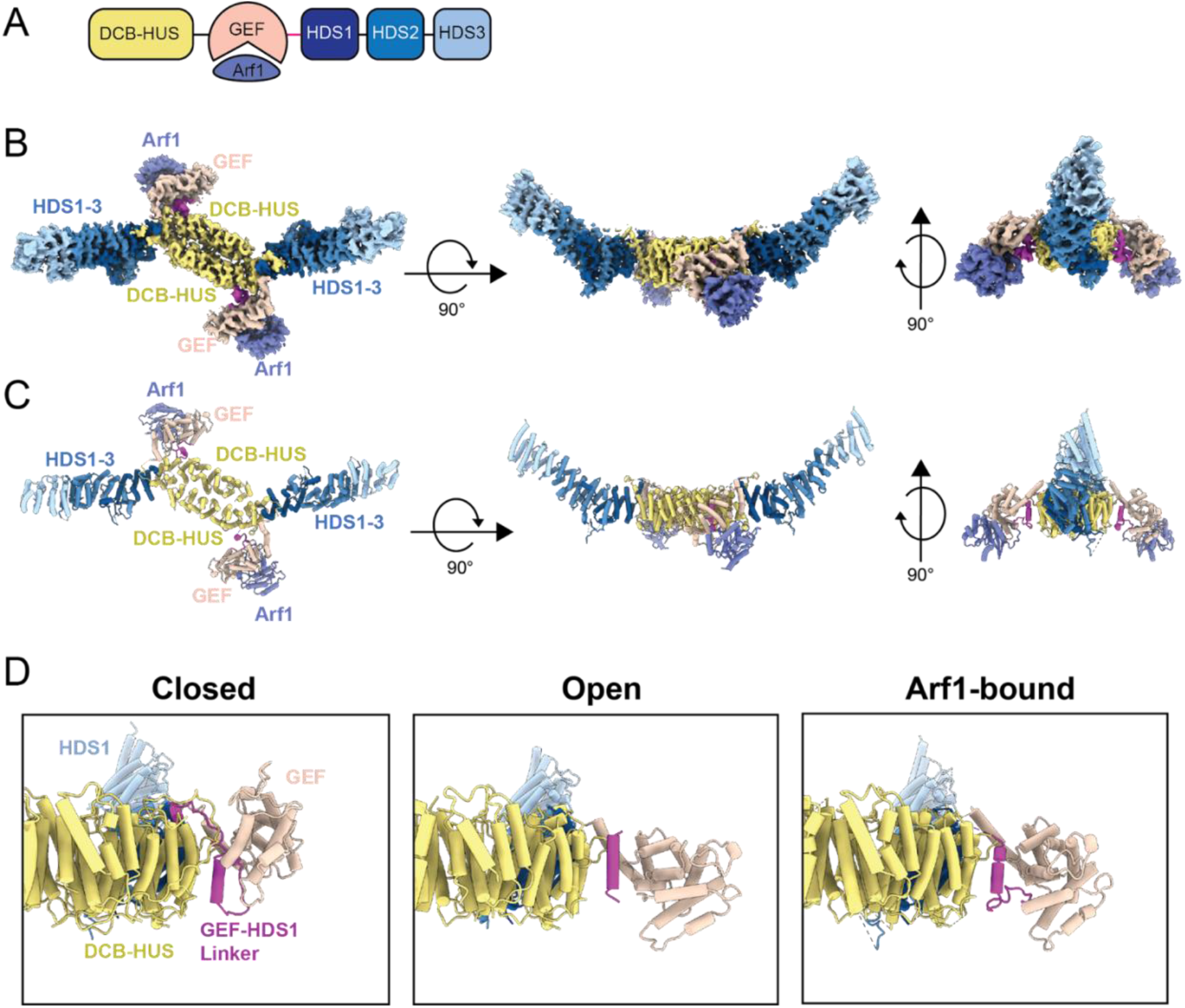
CryoEM structure of a Gea2-Arf1 activation intermediate complex. A) Schematic of the Gea2-Arf1 activation intermediate complex used for cryoEM. B) CryoEM density of the Gea2-Arf1 complex, colored and labeled as in Figure 1, with Arf1 colored purple. C) Atomic model of the Gea2-Arf1 complex. D) Views of the Gea2 GEF domain and GEF-HDS1 linker for each of the three conformations adopted by Gea2 in the Gea2 only (closed and open) and Arf1-bound conformations.

The conserved GEF-HDS1 linker adopts distinct conformations when in the closed, open, and Arf1-bound states (Figs. 4D and S5C,D) and therefore appears important for stabilizing each of these states. In the closed conformation the entire GEF-HDS1 linker is ordered, whereas nearly 20 residues (residue numbers 781-798) at the C-terminal end of the GEF-HDS1 linker are disordered in the open and Arf1-bound structures. To understand why the closed conformation was not observed in the Arf1-bound complexes, we generated a series of models representing different stages of the established Arf1 activation pathway (Fig. 5A-H). To model nucleotide-free Arf1 bound to the closed conformation of Gea2, we superimposed our structure of nucleotide-free Arf1 bound to the GEF domain onto the GEF domain of the closed complex (Fig. 5D). This modeled complex resulted in a steric clash between the ‘switch I’ region of Arf1 and the GEF-HDS1 linker of Gea2 (Fig. 5G). This indicates that the Gea2 closed conformation is incompatible with binding to the nucleotide-free state of Arf1. This steric clash with the closed conformation also explains why the nucleotide-free Gea2-Arf1 activation intermediate adopts an open conformation.

**Figure 5.**
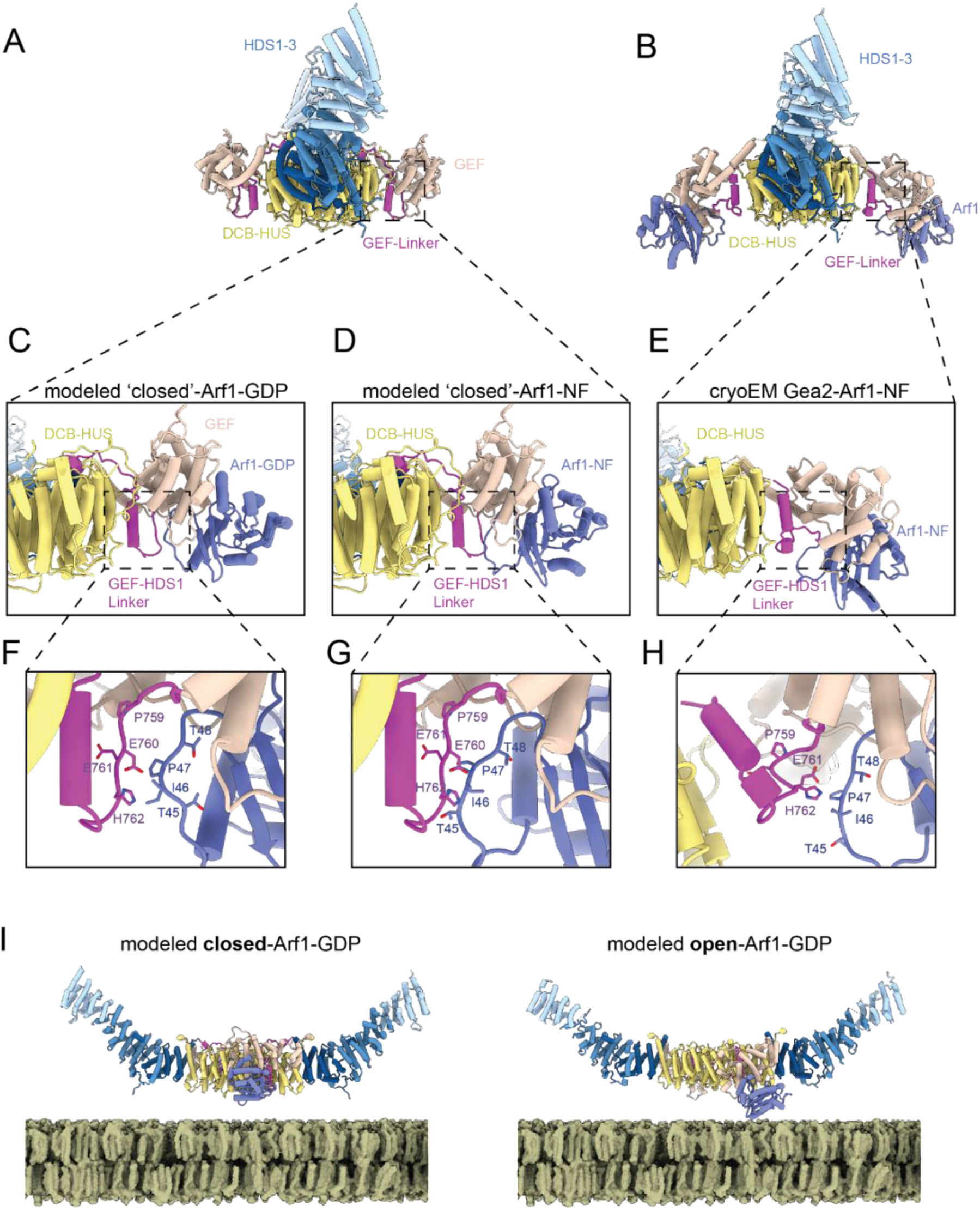
Steric constraints appear to enforce Gea2 conformational change. A) Structure of the closed/closed Gea2 dimer shown for context. B) Structure of the Gea2-Arf1 complex shown for context. C) Close-up view of the modeled Gea2 closed-Arf1-GDP complex. D) Close-up view of the modeled Gea2 closed-Arf1-NF (nucleotide-free) complex. E) Close-up view of the Gea2-Arf1-NF cryoEM structure. F-H) Magnified views of (C-E). Note the steric clash between Arf1 and the GEF-HDS1 linker in (D) and (G). I) Comparison of the modeled closed/closed Gea2-Arf1-GDP complex with the modeled open/open Gea2-Arf1-GDP complex. Note how in the closed conformation, the GEF domain appears more readily able to encounter freely-diffusing Arf1-GDP, compared to the open conformation.

### Evidence for GEF conformational switching during Arf1 nucleotide exchange

These findings raised the question of whether the closed conformation served any role in the nucleotide exchange reaction. We therefore superimposed the published structure of Arf1-GDP bound to the GEF domain from ARNO (Renault et al., 2003) onto the closed conformation of Gea2 (Fig. 5C). In contrast to the nucleotide-free state, Arf1-GDP appears able to bind to Gea2 in the closed conformation without clashes (Fig. 5F), because the configuration of the Arf1 ‘switch I’ region is different in the GDP-bound and nucleotide-free states. This suggests that the closed conformation of Gea2 is compatible with binding to Arf1-GDP.

We were initially puzzled by our observation that the ‘open’ position of the GEF domain in the nucleotide-free Gea2-Arf1 complex appears unsuitable for the initial association event between Gea2 and Arf1-GDP, assuming Gea2 is already membrane-bound. The orientation of the GEF domain active site facing towards the membrane suggested its close proximity to the membrane would preclude it from productively encountering its substrate Arf1-GDP via diffusion, either from the cytosol or along the membrane surface. In contrast, the closed conformation, in which the GEF domain active site is oriented orthogonal to the membrane surface, appears much more suitable for productive encounters with the Arf1-GDP substrate via diffusion than does the open conformation (Fig. 5I).

Taken together, our structural analysis suggests that initial binding to Arf1-GDP likely occurs with the Gea2 GEF domain in the closed conformation (Fig. 5C, F). Subsequent release of GDP, triggered by interaction with the GEF domain, causes Arf1 to adopt its nucleotide-free structure. As this conformation of Arf1 is incompatible with the Gea2 closed state (Fig. 5D, G), the GEF domain likely switches to the open state concurrent with nucleotide release, adopting the nucleotide-free Arf1-bound conformation we observed by cryoEM (Fig. 5E, H). Given the apparent independence of each GEF domain in the dimer, it is also possible that only one GEF domain is able to adopt the open conformation at a time when Gea2 is bound to the membrane.

### A model for activation-coupled membrane insertion of Arf1

When bound to Gea2 in its nucleotide-free state, Arf1 is positioned such that its N-terminus is oriented towards the membrane surface, and we predict it to be in close proximity to the lipid headgroups (Fig. 5I). Although not present in the construct we used to determine the structure of the complex, the N-terminus of Arf1 folds into a membrane-inserting amphipathic helix upon GTP binding (Antonny et al., 1997; Liu et al., 2010). The conformation of Gea2 when bound to the nucleotide-free intermediate therefore appears to prime Arf1 for membrane insertion: GTP binding to the nucleotide-free intermediate induces formation of the N-terminal Arf1 amphipathic helix in a position optimal for its insertion into the cytoplasmic leaflet of the Golgi membrane.

Our structural results and analyses lead us to a complete model for nucleotide exchange-coupled membrane insertion of Arf1 by Gea2 (Fig. 6, Movie S1). Arf1-GDP initially encounters membrane-bound Gea2 in its closed conformation (Fig. 6A-C). Nucleotide release then leads to an open conformation to avoid steric clash with the GEF-HDS1 linker. The resulting open conformation positions the N-terminus of Arf1 optimally for membrane insertion (Fig. 6D). Finally, GTP binding triggers membrane insertion of Arf1 via folding of its myristoylated amphipathic helix and release from Gea2 (Fig. 6E).

**Figure 6.**
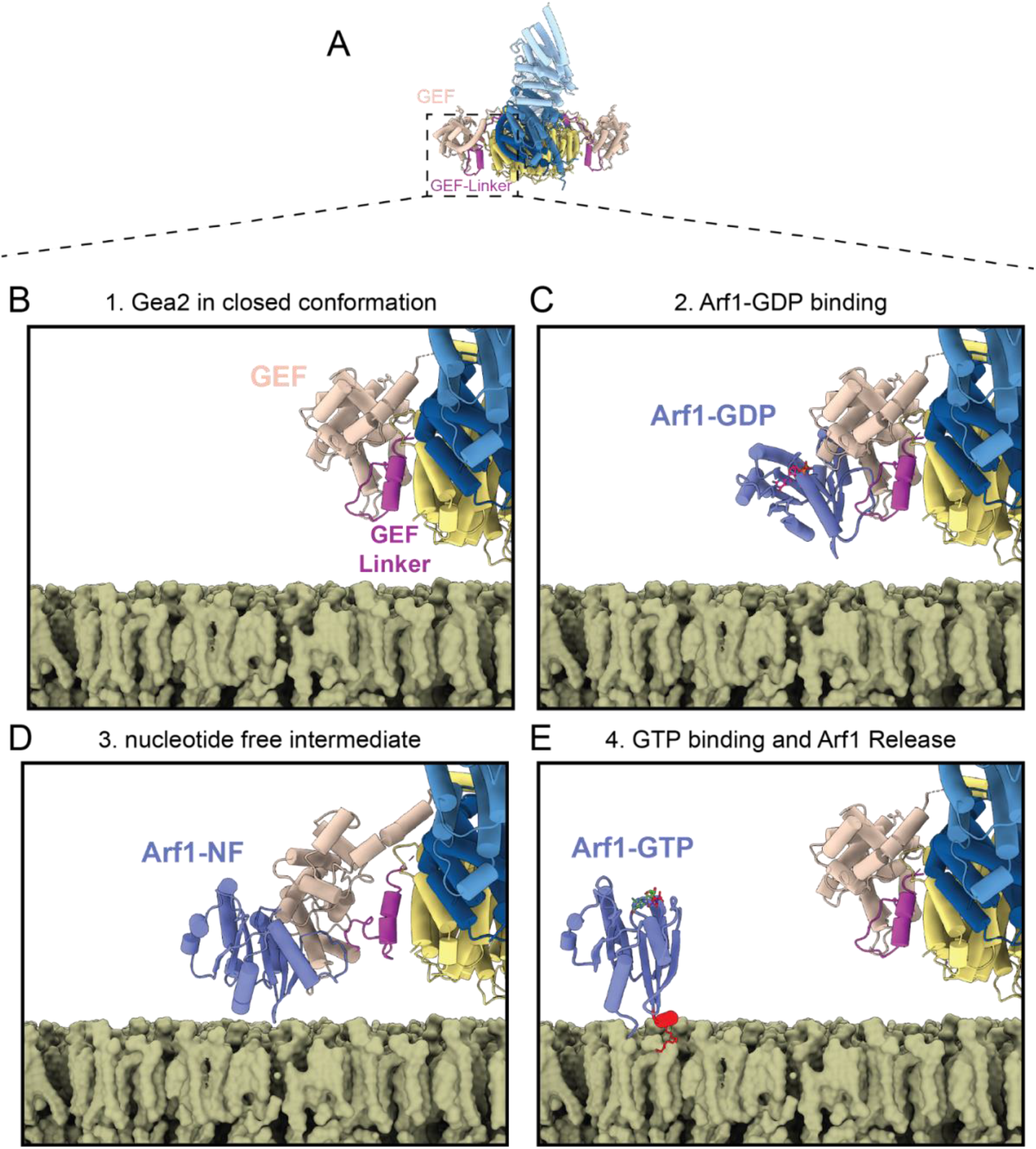
Model for activation of Arf1 by Gea2 on the Golgi membrane surface. A) Gea2 in the closed/closed conformation shown for context. B) In Step 1, at least one of the Gea2 monomers adopt the closed conformation while bound to the membrane surface (the cryoEM structure of one side of the closed/closed conformation shown on a modeled membrane). C) In Step 2, Arf1-GDP binds to the GEF domain (the modeled closed-Arf1-GDP complex is shown). D) In Step 3, GDP dissociates from Arf1 (Arf1-NF = nucleotide free), and the resulting conformation change in Arf1 causes the GEF domain to switch from the closed state to an open state in order to avoid steric clash with Arf1 (the Gea2-Arf1 cryoEM structure is shown). E) In step 4, GTP binding causes another conformation change in Arf1, resulting in folding of its amphipathic helix (colored red) at the membrane surface and dissociation from Gea2 (the NMR structure of Arf1-GTP and cryoEM structure of the closed/closed conformation of Gea2 are shown). The structures of Arf1-GDP and Arf1-GTP were derived from RCSB entries 1R8S (Renault et al., 2003) and 2KSQ (Liu et al., 2010). See also Movie S1.

## Discussion

Arf1 is known for its role as a regulator of the function and regulation of the Golgi complex and recycling endosomes, but its activity has also been implicated in endocytosis, TORC1 kinase signaling, lipid droplet homeostasis, and lysosomal and mitochondrial function (Ackema et al., 2014; Dechant et al., 2014; Kumari and Mayor, 2008; Su et al., 2020; Wilfling et al., 2014). A hallmark of Ras-related “small” GTPases like Arf1 is the structural transitions they undergo during nucleotide exchange and hydrolysis. Arf1 is the founding member of the Arf GTPase family, which includes over 20 proteins in humans which collectively regulate virtually all membrane trafficking pathways (Gillingham and Munro, 2007). Most Arf family GTPases are anchored to the membranes of organelles and vesicles by their N-terminal amphipathic helices. Unlike other Ras-related GTPases, when inactive these membrane-anchoring motifs are masked by direct interaction with the GDP-bound Arf1 nucleotide-binding domain (Amor et al.,1994). In contrast, Rab and Rho family GTPases employ chaperone proteins (guanine-nucleotide displacement inhibitors) to mask their membrane-anchoring motifs in the GDP-bound state (Isomura et al., 1991; Soldati et al., 1994). GTP-binding exposes the Arf amphipathic helix, inducing stable membrane binding (Antonny et al., 1997). Although membrane insertion of GTP-bound Arf proteins is favorable, there is likely a kinetic ‘activation energy’ barrier that slows the membrane-insertion step, as it requires lipids to rearrange in order to accommodate the amphipathic helix. Our structural findings point to a mechanism for how Gea2 may reduce this kinetic barrier by positioning Arf1 optimally for membrane insertion.

To our knowledge, conformational change of a GEF during the nucleotide exchange reaction has not been reported. Several GEFs are known to be autoinhibited and/or allosterically activated, and the structural basis for autoinhibition and activation has been documented for several GEFs, including the Ras-GEF SOS (Gureasko et al., 2008;Sondermann et al., 2004), the Rab-GEF Rabex5 (Delprato and Lambright, 2007; Lauer et al.,2019; Zhang et al., 2014), the Arf-GEF Cytohesin/Grp1 (Das et al., 2019; DiNitto et al., 2007;Malaby et al., 2013), and the Rho-GEF Vav (Yu et al., 2010). In the context of autoinhibition and allosteric activation, GEF conformational change is usually coupled to phosphorylation or binding to a regulatory protein or lipid and is a prerequisite for the nucleotide exchange reaction. In contrast, Gea2 appears to capitalize on the conformational changes its substrate GTPase undergoes during nucleotide exchange to drive its own conformational change during the activation reaction.

A mutation has been identified in geaA, the *Aspergillus nidulans* homolog of *S. cerevisiae* Gea2, corresponding to a Y1001C substitution in Gea2, that partially suppressed the loss of the *A. nidulans* homolog of Sec7, hypB (Arst et al., 2014). Remarkably, this Y-to-C substitution mutation shifted the localization of geaA from early-Golgi compartments towards later Golgi compartments normally occupied by hypB. Our findings provide a mechanistic interpretation of this observation, as we have identified Y1001 as a critical residue for Gea2 membrane interaction through our use of the Y1001D mutant. An interesting possibility is that the Y-to-C substitution, by modulating but not eliminating the hydrophobicity of the amphipathic helix, alters which membranes are most favored for stable binding due to their compositions or biophysical properties. We note that in contrast to the results reported for *A. nidulans* geaA, we found that the equivalent Y-to-C substitution did not enable Gea2 to suppress loss of Sec7 in *S. cerevisiae* (Gustafson, 2017). This highlights the proposed roles of regulatory protein-protein interactions in directing the localization of the Golgi Arf-GEFs to specific compartments (Christis and Munro, 2012; Gustafson and Fromme, 2017; Lowery et al., 2013; McDonold and Fromme,2014; Monetta et al., 2007; Richardson et al., 2012).

There are likely to be both similarities among and differences between the structural mechanisms underlying Arf1 activation by Gea2 and Sec7. Previous work on Sec7 highlighted the influence of the DCB-HUS domain on the activity of the GEF domain for activation of Arf1 on the membrane surface (Halaby and Fromme, 2018; Richardson et al., 2016). However, Sec7 likely adopts a very different overall architecture because Sec7 dimerizes via its HDS4 domain (Richardson et al., 2016). Sec7 is also regulated by distinct positive-feedback, autoinhibition, and cross-talk mechanisms (McDonold and Fromme, 2014; Richardson et al., 2012) and prefers more anionic membranes compared to Gea1/2 (Gustafson and Fromme, 2017).

Although we have now identified how Gea2 interacts with membranes, it remains unresolved how it achieves its specific localization. Both Gea1 and Gea2, as well as GBF1, interact with Rab1/Ypt1, which likely recruit these Arf-GEFs to the Golgi, yet Gea1 and Gea2 localize to distinct Golgi compartments (Gustafson and Fromme, 2017; Monetta et al., 2007). Future studies are required to characterize the Gea2-Ypt1 interaction and determine how Gea1 and Gea2 achieve their specific localization.

## Methods

### Protein purifications and Gea2-Arf1 complex formation

Full-length *S. cerevisiae* Gea2 was cloned with an N-terminal cleavable 6xHis-tag into the pPICZ vector (Table S1), then purified using *Pichia pastoris*. An overnight culture of “BMGY” media was used to inoculate a 200mL BMGY starter culture. After 8 hours of shaking at 30°C, 120 mL of this starter culture was used to inoculate 6 liters of “autoinduction media” (Lee et al.,2017) and then shaken overnight at 30°C. After overnight growth, additional methanol was added (equivalent to additional 0.5% final concentration) and the cultures were shaken for an additional 24 hours at 30°C. Cells were harvested by centrifugation (2000 g, 10 min), resuspended in lysis buffer (50 mM Tris pH 8.0, 500 mM NaCl, 10% glycerol, 20 mM imidazole 10 mM βME), and lysed under liquid nitrogen using a SPEX 6875D freezer mill. Lysed cells were cleared using centrifugation (40,000 g, 1 hr) and the supernatant was incubated with 1 mL Ni^2+^-NTA resin for 1 hr. Resin was washed with lysis buffer and the protein was eluted with elution buffer (50 mM Tris pH 8.0, 500 mM NaCl, 10% glycerol, 500 mM imidazole, 10 mM BME). The elute was then diluted 5x with Buffer A (20 mM Tris pH 8.0, 1 mM DTT) and subjected to ion exchange using a MonoQ column (Buffer B = Buffer A + 1 M NaCl). Fractions were visualized by SDS page and pooled fractions were concentrated to 500 μL total volume then treated with 50 μL of 1 mg/mL TEV protease overnight at 4°C. The sample was further purified by size exclusion chromatography using a Superdex 200 Increase column equilibrated in SEC buffer (20 mM Tris pH 8.0, 150 mM NaCl, 1 mM DTT). The Y1001D mutant was purified using the same procedure.

*S. cerevisiae* ΔN17-Arf1 and myristoylated-Arf1 were purified as previously described (Richardson and Fromme, 2015; Richardson et al., 2012).

The Gea2-Arf1 complex was prepared by incubating 1 mg of Gea2, 5 mg ΔN17Arf1, and 250 units alkaline phosphatase in 1.5 ml reaction volume at 4°C overnight. The complex was then purified by size exclusion chromatography using a Superdex 200 Increase column equilibrated in SEC buffer (20 mM Tris pH 8.0, 300 mM NaCl, 1 mM DTT).

### CryoEM Sample preparation and data collection

3.5 μL of Gea2 or the Gea2-Arf1 complex, at ~5 mg/mL in SEC Buffer containing 2 mM fluorinated fos-choline-8 (Anatrace, cat# F300F), was applied to glow discharged Quantifoil R1.2/1.3 grids, blotted for 5 seconds, then plunge-frozen into liquid ethane using a Vitrobot Mark IV. Imaging was done at 63kX nominal magnification on a Talos Arctica operating at 200kV equipped with a K3 detector and BioQuantum energy filter. For Gea2 alone, ~8000 movies were collected over multiple sessions, and for the Gea2-Arf1 complex ~2500 movies were collected. Movie exposures were collected using SerialEM (Mastronarde, 2005) using the multi-shot feature with coma correction. All data was collected using 100 frames per movie exposure with a total dose of ~50 e^-^/ Å^2^.

### CryoEM data processing

#### Gea2 alone

Movie exposures were motion-corrected and dose-corrected using MotionCor2 (Zheng et al., 2017). Corrected micrographs were imported into cryoSPARC (Punjani et al., 2017) and then subjected to patch-CTF estimation. Particle picking was performed via TOPAZ (Bepler et al., 2019, 2020) using a ‘general’ model. Picked particles were parsed with 2D classification and rounds of 3D classification (see Fig S1). A clean particle stack was generated and imported into RELION 3.1 (Zivanov et al., 2018, 2020) and particles were 3D classified revealing three distinct conformations. Particles in each of the these three major classes were kept separate for the rest of the processing steps. Particles were subjected to multiple rounds of CTF refinement and Bayesian polishing (Zivanov et al., 2019). C2 symmetry was enforced during refinements of the open and closed states. After the iterative refinement process converged, particles from the closed/closed and open/open states were symmetry expanded and signal subtracted using a monomer mask (Nakane et al., 2018). For the closed/open state, an additional refinement was performed with C2 symmetry enforced in order to perform symmetry expansion and monomer particle subtraction. 3D classification was then used to generate separated particle stacks for the open and closed monomers. Following monomer refinements, subsequent signal subtraction and local refinements were performed separately on the N and C terminal regions. An additional signal subtraction and focused refinement was performed for the dimer interface of each of the three states (open, closed, and hemi). Density modification (Terwilliger et al.,2020) was then used to further improve all of the focused maps. Composite maps used for model building and refinement of each of the three dimeric conformations were generated with ‘Combine Focused Maps’ in Phenix (Liebschner et al., 2019). See Figs. S2 and S3, and Table 1.

#### Gea2-Arf1 complex

The cryoEM data collected for the Gea2-Arf1 complex was processed using the same procedure described above for Gea2 alone. 3D classification indicated that the sample was conformationally homogeneous, adopting a single conformation. After symmetry expansion and signal subtraction, focused refinements were performed on the DCB-HUS, GEF, and HDS1-3 regions. Density modification (Terwilliger et al., 2020) was used to further improve all of the focused maps, and composite maps used for model building and refinement were generated with ‘Combine Focused Maps’ in Phenix (Liebschner et al., 2019). See Fig. S6 and Table 1.

### Atomic model building and refinement

The composite maps described above were used for atomic model building and refinement. Model building in Coot (Emsley et al., 2010) was guided by the AlphaFold prediction of Gea2 (Jumper et al., 2021) and by the Gea2 GEF domain - Arf1 crystal structure (Goldberg, 1998). Real space refinement and model validation was carried out using Phenix (Afonine et al., 2018; Emsley et al., 2010). See Figs. S3 and S6 and Table 1.

### Yeast complementation assay

Gea2-expressing yeast plasmids (Table S1) were transformed into a Gea1/2 yeast shuffling strain (*gea1*Δ *gea2*Δ strain CFY2872, Table S2) and grown overnight at 30°C. Cultures were normalized by OD600 and serial diluted on selection media. Plates were then incubated for three days at 30°C before imaging.

### Fluorescence microscopy

Gea2-expressing yeast plasmids (Table S1) were transformed into *gea2*Δ yeast strain (CFY1470, Table S2) and grown at 30°C in selection media to an OD600 of 0.6. Cells were added to an imaging dish (MatTek), allowed to settle for 10 mins, then washed with fresh media. Cells were imaged using a CSU-X spinning-disk confocal system (Intelligent Imaging Innovations) with a DMI6000 B microscope (Leica), 100X 1.46 NA oil immersion objective, and a QuantME EMCCD camera (Photmetrics), using with a 200 μs exposure time.

### Liposome Preparation

Liposomes were prepared in HK buffer (20mM HEPES pH 7.5, 150 mM KOAc) as described previously (Richardson and Fromme, 2015). Hydrated lipid mixes were extruded with 100 nm filters for GEF assays and 400 nm filters for the membrane-binding liposome pelleting assay.

### In vitro membrane-binding assay

Liposome pelleting assays were performed as previously described (Gustafson and Fromme,2017; Paczkowski and Fromme, 2016), using liposomes composed of 94% DOPC, 5% Nickel-DOGS, and 1% DiR lipids. 500 μg liposomes were incubated with 8 ug of protein in 50 μL total reaction volume in HK buffer for 10 mins at room temperature. Reactions were then subjected to ultracentrifugation (128,000g for 10 mins). The supernatant was separated and the liposome pellet was resuspended in HK buffer. Supernatant and pellet samples were analyzed by SDS-PAGE.

### In vitro GEF activity assay

GEF activity assays were performed as previously described, monitoring native tryptophan fluorescence (Gustafson and Fromme, 2017; Richardson and Fromme, 2015) using 99% DOPC and 1% DiR lipids. All reactions were performed in HKM buffer (20mM HEPES pH 7.5, 150 mM KOAc, 1 mM MgCl_2_) at 30°C. Myristoylated-Arf1 activation reactions were performed by incubating 333 μM liposomes, 200 nM Gea2, 200 μM GTP for 2 minutes before adding 1 μM myr-Arf1. ΔN-Arf1 activation was assessed in similar reactions, except liposomes were omitted, Gea2 concentration was 25 nM, and ΔN-Arf1 concentration was 500 nM.

## Data Availability

CryoEM maps and atomic coordinates will be available from the EMDB and RCSB upon publication.

## Acknowledgements

We acknowledge the Cornell Center for Materials Research (CCMR), especially K. Spoth and M. Silvestry-Ramos, for access and support of electron microscopy sample preparation and data collection. We thank members of the Fromme lab for helpful advice and discussions. We thank the Accardi lab for suggesting the use of fos-choline-8. This study was funded by NIH grant R35GM136258 to J.C.F. The CCMR is supported by NSF grant DMR-1719875.

## Author contributions

AJM performed all experiments and data analysis. MAG performed extensive protein crystallization trials and optimization. JCF supervised the project and obtained funding. AJM and JCF wrote the manuscript with input from MAG.

## Competing interests

The authors declare no competing interests.

## Supplementary Materials

**Figures S1–S7**

**Tables S1–S2**

**Movie S1**

**Figure S1.**
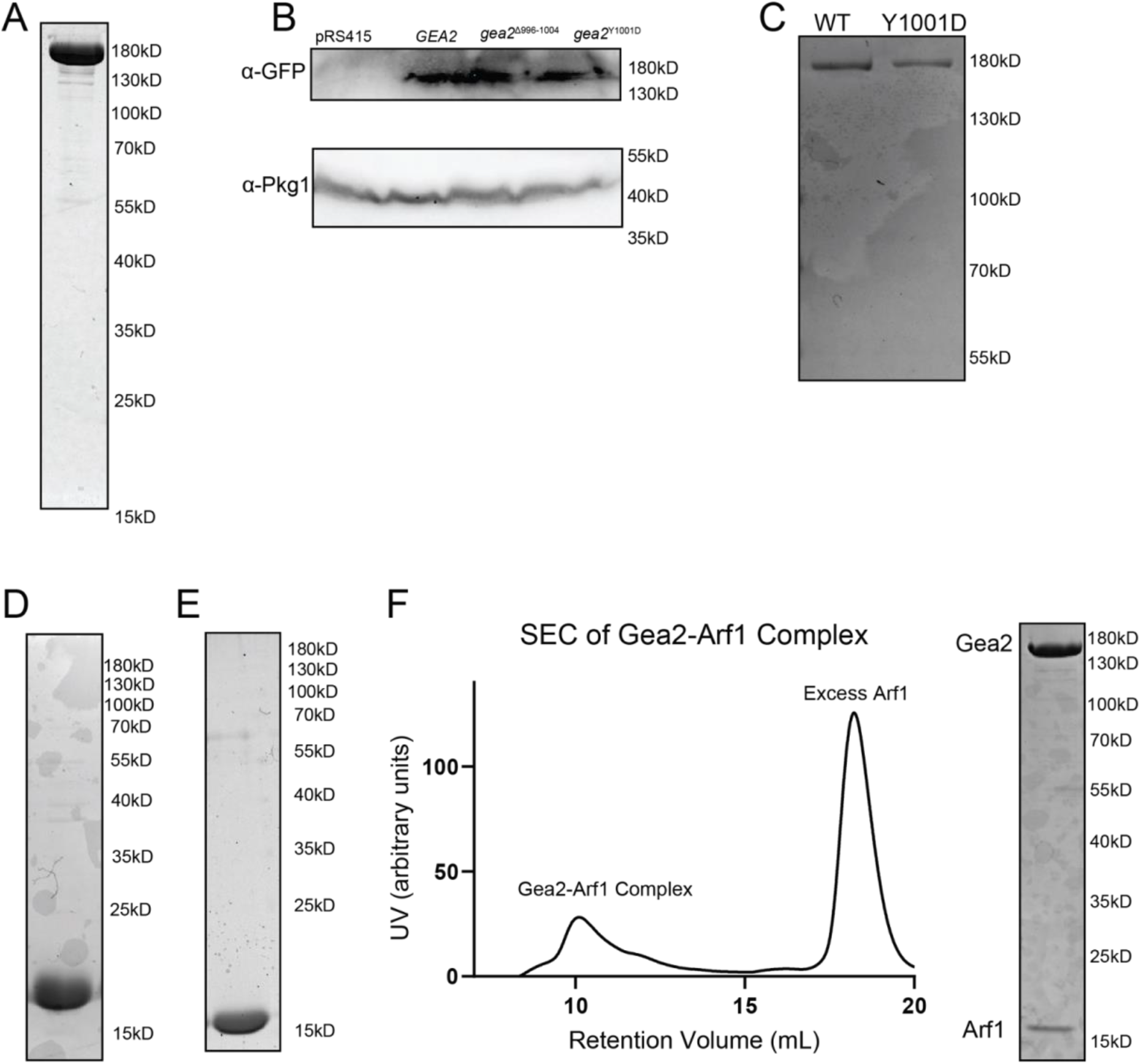
Protein reagents used in this study. A) SDS-PAGE analysis of *S. cerevisiae* Gea2 purified from *P. pastoris*. B) Immunoblot showing expression of Gea2-GFP constructs used in yeast localization experiments. C) SDS-PAGE analysis of wild-type and Y1001D mutant *S. cerevisiae* Gea2 proteins purified from *P. pastoris*. D) SDS-PAGE analysis of Myristoylated, full-length *S. cerevisiae* Arf1 purified from *E. coli*. E) SDS-PAGE analysis of ΔN17-mutant *S. cerevisiae* Arf1 purified from *E. coli*. F) Gel filtration chromatography trace and SDS-PAGE analysis of the Gea2-Arf1 activation intermediate complex.

**Figure S2.**
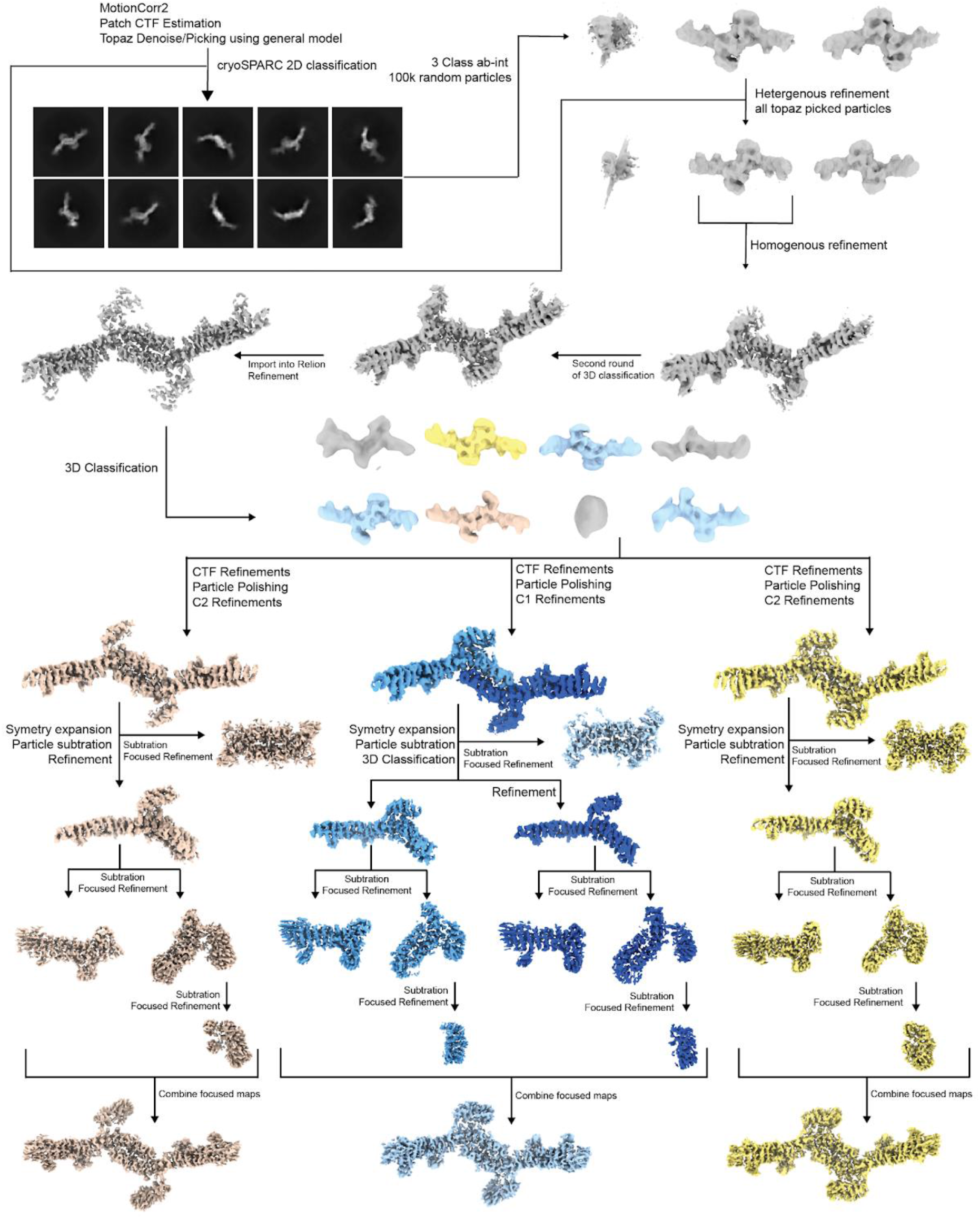
Gea2 cryoEM data processing. Flowchart illustrating the data processing strategy for the Gea2 cryoEM data (see Methods).

**Figure S3.**
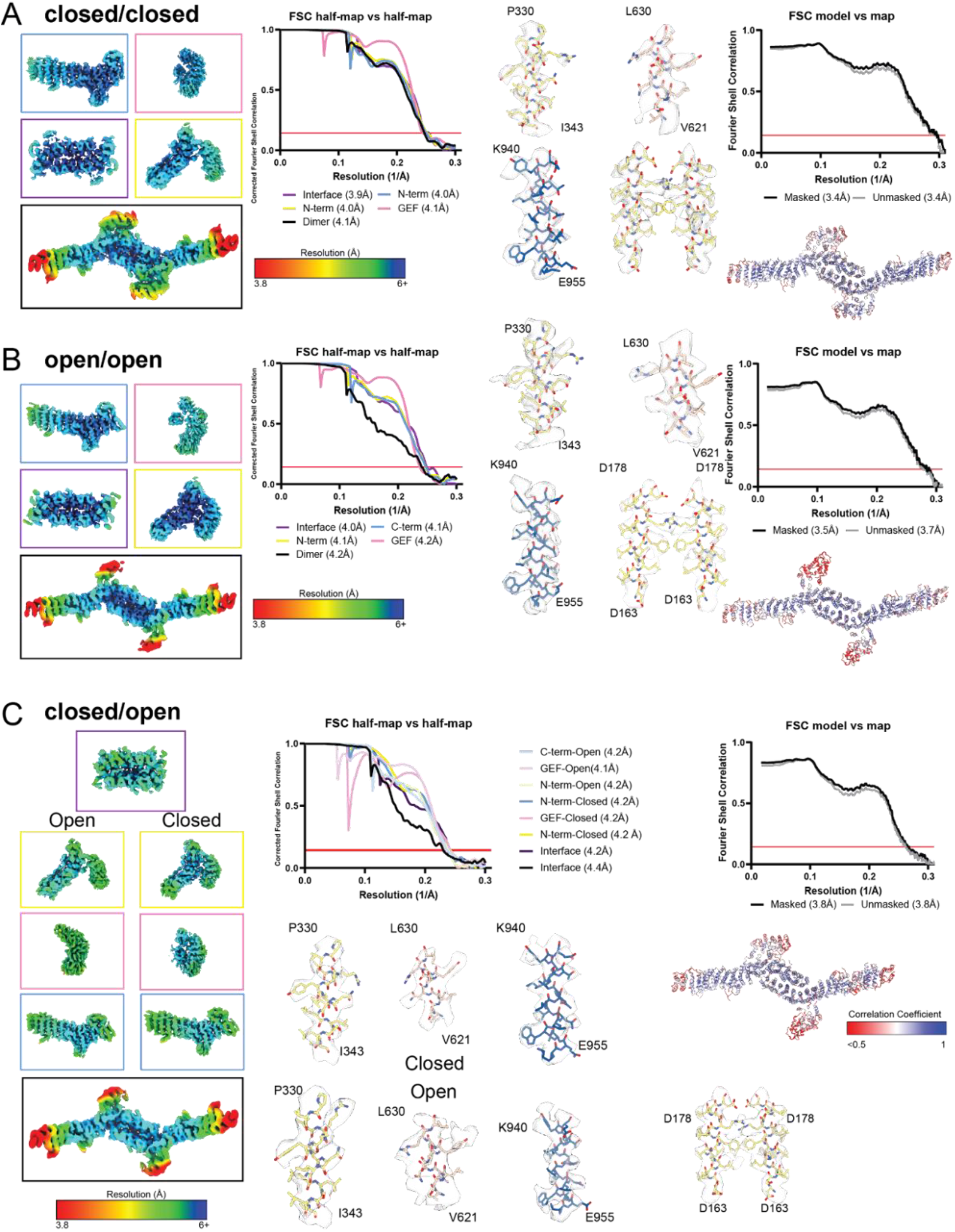
Gea2 cryoEM map and model validation. A) Fourier shell correlation plots and example cryoEM density for focused refinements are shown for the cryoEM map and model of the Gea2 closed/closed conformation. B) Same but for the open/open conformation. C) Same but for the closed/open conformation. Note that the open conformation of the GEF domain exhibits more flexibility compared to the closed conformation.

**Figure S4.**
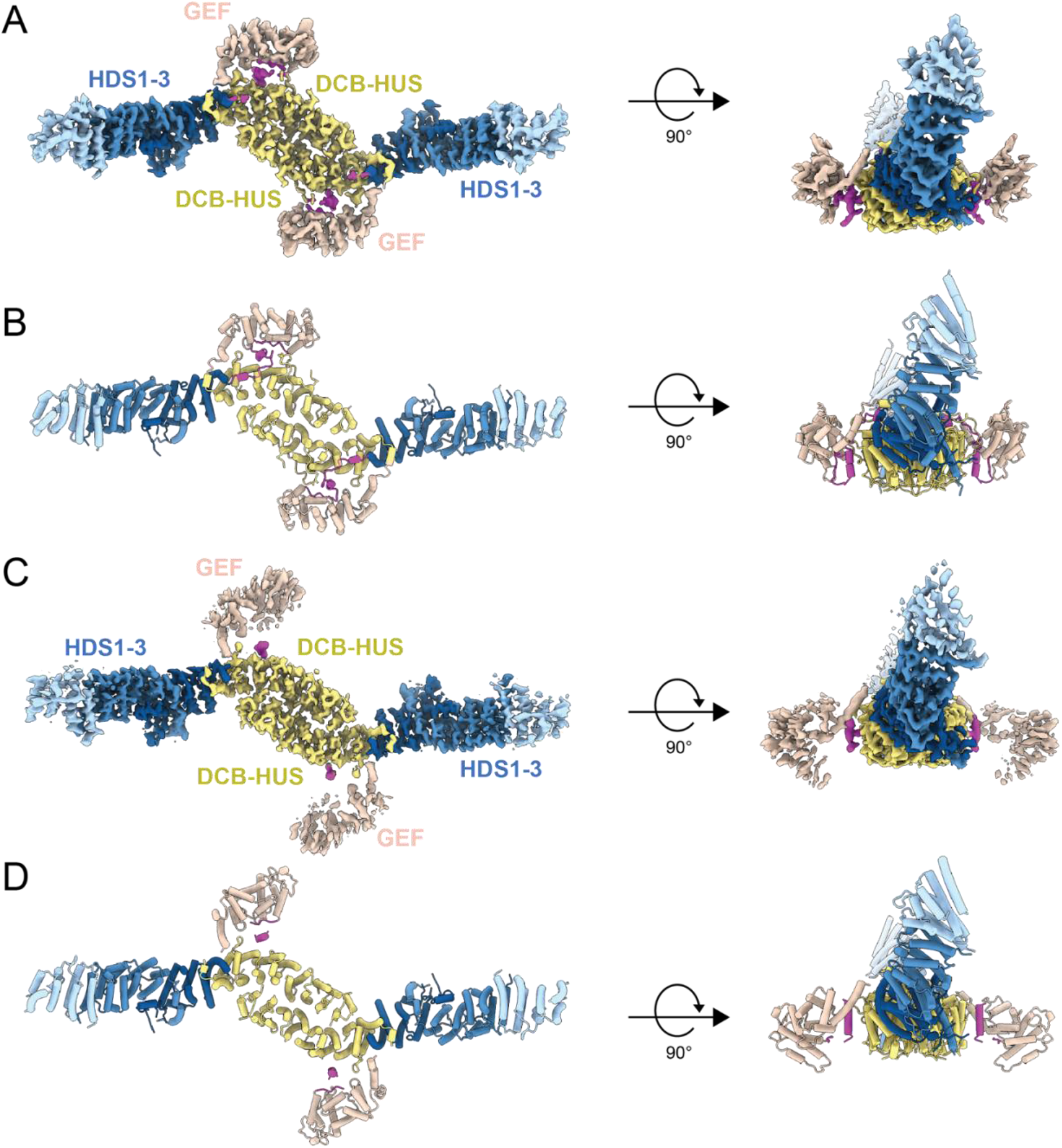
The closed/closed and open/open conformations of Gea2. A) CryoEM density of the Gea2 dimer in its closed/closed conformation. Coloring as in Fig. 1, the GEF-HDS1 linker is colored magenta. B) Atomic model of the Gea2 closed/closed dimer, shown in cartoon depiction. C) CryoEM density of the Gea2 dimer in its open/open conformation. Coloring as in Fig. 1, the GEF-HDS1 linker is colored magenta. D) Atomic model of the Gea2 open/open dimer, shown in cartoon depiction.

**Figure S5.**
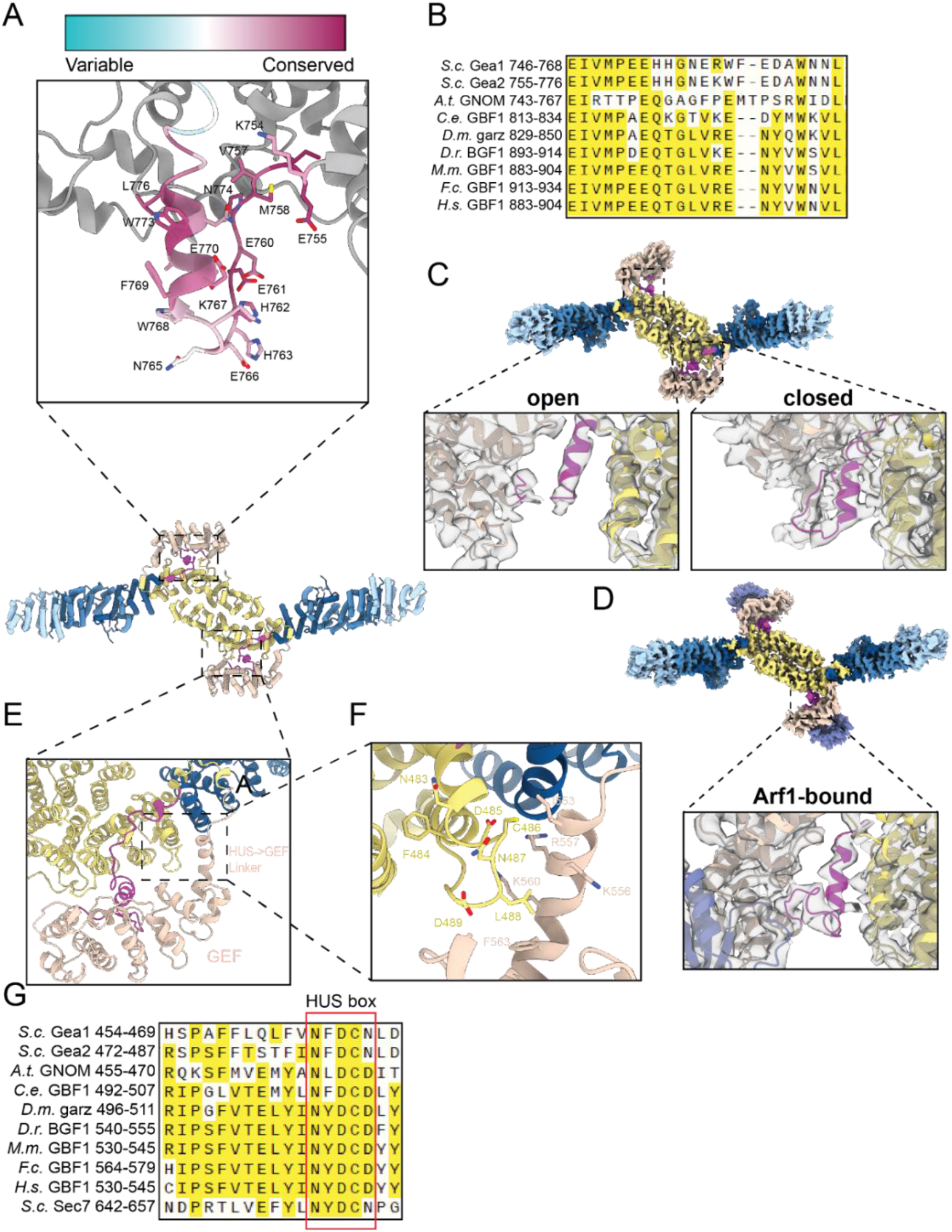
Ordered linkers connect the GEF domain to the HUS and HDS1 domains. A) Close-up view of the GEF-HDS1 linker in the closed monomer colored by conservation as indicated. B) Sequence alignment of the GEF-HDS1 linker from Gea2 homologs in several model organisms and humans. Yellow highlights indicate identical residues at a given position. C) Views of the GEF-HDS1 linker cryoEM density in the open (left) and closed (right) monomers. D) View of the GEF-HDS1 linker cryoEM density in the Gea2-Arf1 complex. E,F) Close-up views of the HUS-GEF linker and ‘HUS-box’ region in the closed monomer. G) Sequence alignment of the HUS-GEF linker from Gea2 homologs in several models organisms and humans. Yellow highlights indicate identical residues at a given position.

**Figure S6.**
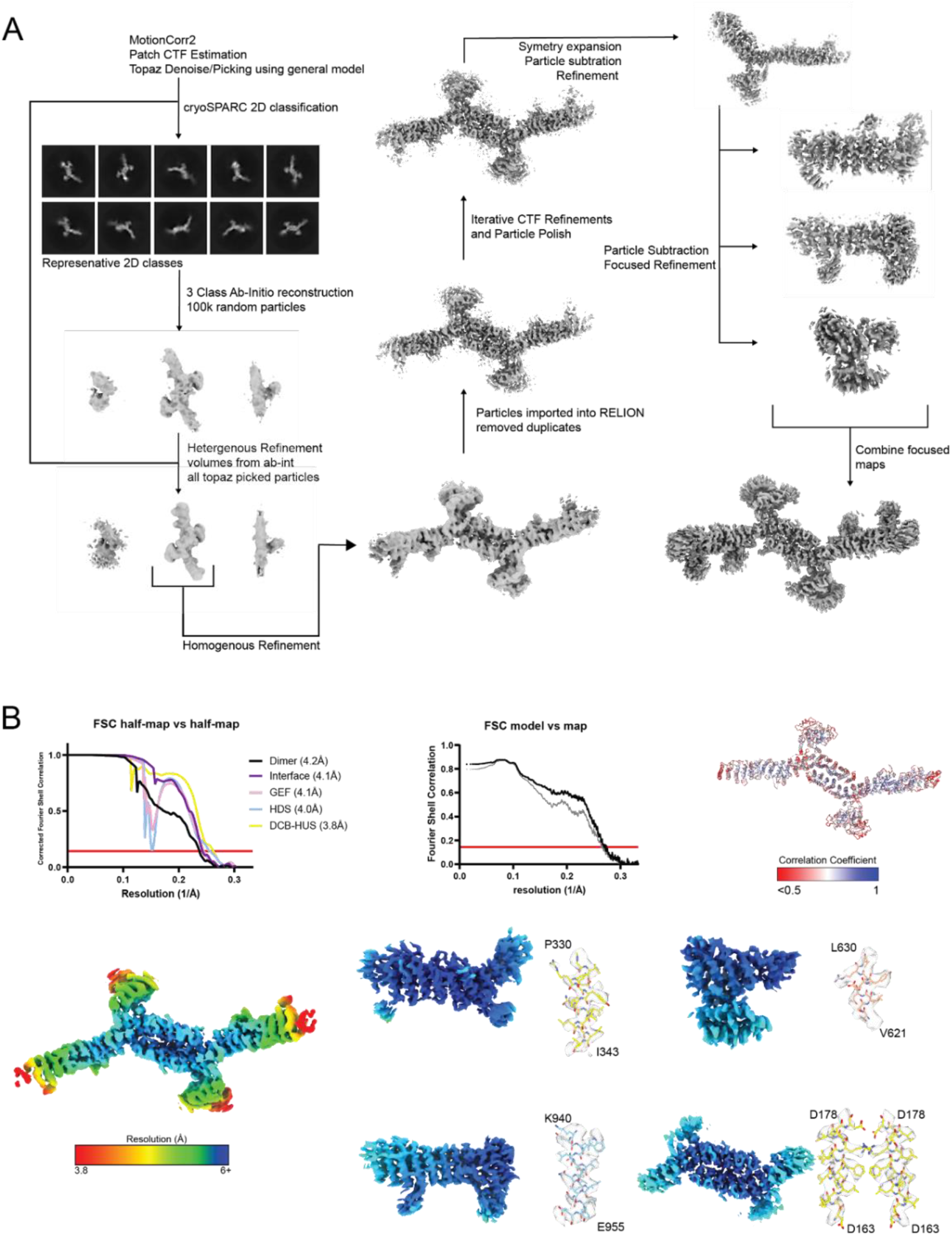
Gea2-Arf1 activation intermediate complex cryoEM data processing. A) Flowchart illustrating the data processing workflow for the Gea2-Arf1 complex cryoEM data (see Methods). B) Fourier shell correlation plots and example cryoEM density for focused refinements are shown for the cryoEM map and model of the Gea2-Arf1 complex.

**Figure S7.**
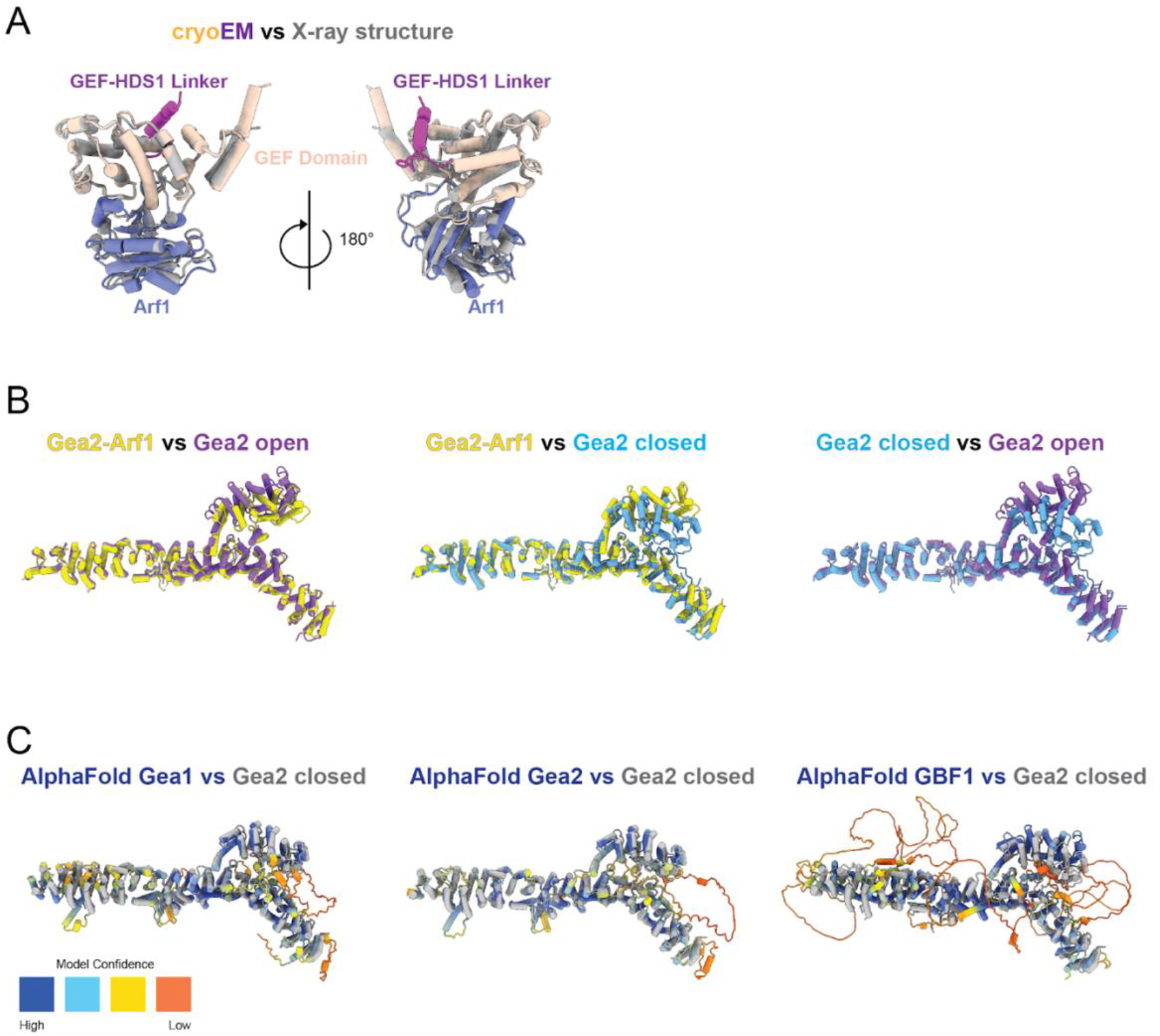
Structural comparisons. A) CryoEM structure of the Gea2 GEF domain bound to nucleotide-free Arf1 superimposed on the crystal structure of the Gea2 GEF domain bound to nucleotide-free Arf1 (Goldberg, 1998). B) Pairwise superpositions of the Gea2 open, closed, and Arf1-bound monomers. C) Superpositions of the Gea2 closed monomer onto AlphaFold models, colored by prediction confidence, of *S. cerevisiae* Gea1, *S. cerevisiae* Gea2, and human GBF1 (Jumper et al., 2021).

**Movie S1. Structural model for Arf1 activation by Gea2 on a membrane surface.**

At the beginning of the movie, a close-up of a single GEF domain of Gea2 is shown in the closed conformation. Then Arf1-GDP, which is freely diffusing in the cytoplasm, encounters and binds to the GEF domain. GDP then dissociates and the conformation of Arf1 changes to adopt its nucleotide-free state. At the same time, the conformation of Gea2 changes to an open-state, resulting in the Gea2-Arf1 complex structure observed by cryoEM. GTP then binds to Arf1, triggering a change in Arf1 conformation and dissociation from Gea2. Upon adopting its active state, the N-terminal amphipathic helix of Arf1 folds and inserts into the cytoplasmic leaflet of the organelle membrane. The structures of Arf1-GDP and Arf1-GTP were derived from RCSB entries 1R8S (Renault et al., 2003) and 2KSQ (Liu et al., 2010). See also Figure 6.

**Table S1.**
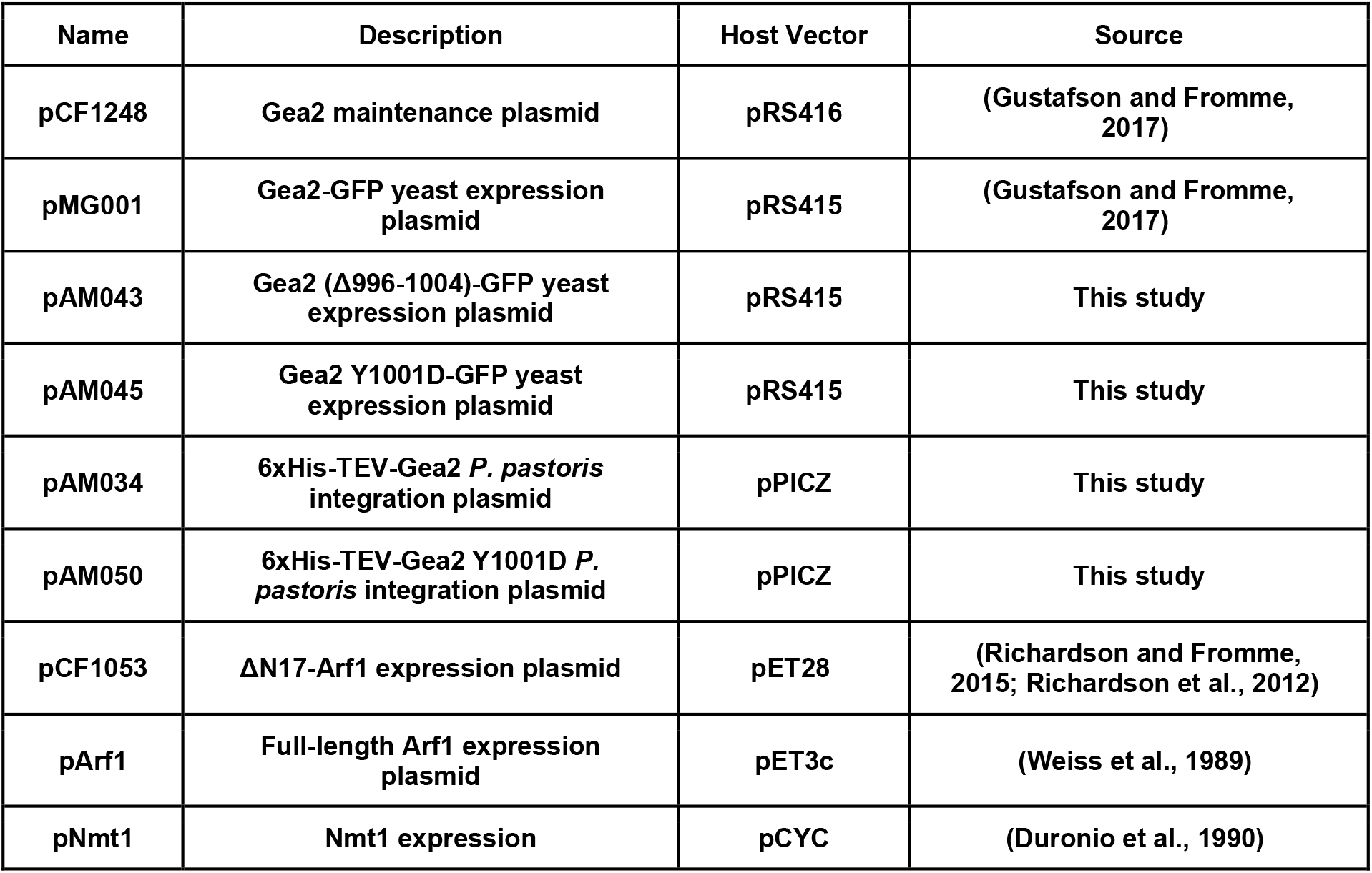
Plasmids.

| Name | Description | Host Vector | Source |
| --- | --- | --- | --- |
| pCF1248 | Gea2 maintenance plasmid | pRS416 | (Gustafson and Fromme, 2017) |
| pMG001 | Gea2-GFP yeast expression plasmid | pRS415 | (Gustafson and Fromme, 2017) |
| pAM043 | Gea2 (Δ996-1004)-GFP yeast expression plasmid | pRS415 | This study |
| pAM045 | Gea2 Y1001D-GFP yeast expression plasmid | pRS415 | This study |
| pAM034 | 6xHis-TEV-Gea2 *P. pastoris* integration plasmid | pPICZ | This study |
| pAM050 | 6xHis-TEV-Gea2 Y1001D *P. pastoris* integration plasmid | pPICZ | This study |
| pCF1053 | ΔN17-Arf1 expression plasmid | pET28 | (Richardson and Fromme, 2015; Richardson et al., 2012) |
| pArf1 | Full-length Arf1 expression plasmid | pET3c | (Weiss et al., 1989) |
| pNmt1 | Nmt1 expression | pCYC | (Duronio et al., 1990) |

**Table S2.** Strains.

| **Name** | **Species** | **Genotype** | **Source** |
| --- | --- | --- | --- |
| SEY6210 | *S. cerevisiae* | *MATα suc2-Δ9 ura3-52 his3-Δ200 leu2-3,112 lys2-801 trp1-Δ901* | (Robinson et al., 1988) |
| CFY2872 | *S. cerevisiae* | *BY4741α gea1Δ::KanMX gea2Δ::HIS3 +pCF1248* | (Gustafson and Fromme, 2017; Liu et al., 2010) |
| CFY1470 | *S. cerevisiae* | SEY6210 *gea2Δ::KanMX* | This study |
| CFY3882 | *P. pastoris* | KM71H *pAOX1::6xHis-TEV-Gea2::BleoR* | This study |
| CFY4619 | *P. pastoris* | KM71H *pAOX1::6xHis-TEV-Gea2 (Y100D)::BleoR* | This study |

## Notes

### Competing Interest Statement

The authors have declared no competing interest.

## References

Ackema, K.B., Hench, J., Böckler, S., Wang, S.C., Sauder, U., Mergentaler, H., Westermann, B., Bard, F., Frank, S., and Spang, A. (2014). The small GTPase Arf1 modulates mitochondrial morphology and function. EMBO J. 33, 2659–2675.

Adarska, P., Wong-Dilworth, L., and Bottanelli, F. (2021). ARF GTPases and Their Ubiquitous Role in Intracellular Trafficking Beyond the Golgi. Front Cell Dev Biol 9, 679046.

Afonine, P.V., Klaholz, B.P., Moriarty, N.W., Poon, B.K., Sobolev, O.V., Terwilliger, T.C., Adams, P.D., and Urzhumtsev, A. (2018). New tools for the analysis and validation of cryo-EM maps and atomic models. Acta Crystallogr D Struct Biol 74, 814–840.

Aizel, K., Biou, V., Navaza, J., Duarte, L.V., Campanacci, V., Cherfils, J., and Zeghouf, M. (2013). Integrated conformational and lipid-sensing regulation of endosomal ArfGEF BRAG2. PLoS Biol. 11, e1001652.

Amor, J.C., Harrison, D.H., Kahn, R.A., and Ringe, D. (1994). Structure of the human ADP-ribosylation factor 1 complexed with GDP. Nature 372, 704–708.

Antonny, B., Beraud-Dufour, S., Chardin, P., and Chabre, M. (1997). N-terminal hydrophobic residues of the G-protein ADP-ribosylation factor-1 insert into membrane phospholipids upon GDP to GTP exchange. Biochemistry 36, 4675–4684.

Arst, H.N., Jr, Hernandez-Gonzalez, M., Peñalva, M.A., and Pantazopoulou, A. (2014). GBF/Gea mutant with a single substitution sustains fungal growth in the absence of BIG/Sec7. FEBS Lett. 588, 4799–4806.

Bepler, T., Morin, A., Rapp, M., Brasch, J., Shapiro, L., Noble, A.J., and Berger, B. (2019). Positive-unlabeled convolutional neural networks for particle picking in cryo-electron micrographs. Nat. Methods 16, 1153–1160.

Bepler, T., Kelley, K., Noble, A.J., and Berger, B. (2020). Topaz-Denoise: general deep denoising models for cryoEM and cryoET. Nat. Commun. 11, 5208.

Bhatt, J.M., Viktorova, E.G., Busby, T., Wyrozumska, P., Newman, L.E., Lin, H., Lee, E., Wright, J., Belov, G.A., Kahn, R.A., et al. (2016). Oligomerization of the Sec7 domain Arf guanine nucleotide exchange factor GBF1 is dispensable for Golgi localization and function but regulates degradation. Am. J. Physiol. Cell Physiol. 310, C456–C469.

Bouvet, S., Golinelli-Cohen, M.-P., Contremoulins, V., and Jackson, C.L. (2013). Targeting of the Arf-GEF GBF1 to lipid droplets and Golgi membranes. J. Cell Sci. 126, 4794–4805.

Bui, Q.T., Golinelli-Cohen, M.-P., and Jackson, C.L. (2009). Large Arf1 guanine nucleotide exchange factors: evolution, domain structure, and roles in membrane trafficking and human disease. Mol. Genet. Genomics 282, 329–350.

Casanova, J.E. (2007). Regulation of Arf activation: the Sec7 family of guanine nucleotide exchange factors. Traffic 8, 1476–1485.

Cherfils, J. (2014). Arf GTPases and their effectors: assembling multivalent membrane-binding platforms. Curr. Opin. Struct. Biol. 29, 67–76.

Christis, C., and Munro, S. (2012). The small G protein Arl1 directs the trans-Golgi-specific targeting of the Arf1 exchange factors BIG1 and BIG2. J. Cell Biol. 196, 327–335.

Claude, A., Zhao, B.P., Kuziemsky, C.E., Dahan, S., Berger, S.J., Yan, J.P., Armold, A.D., Sullivan, E.M., and Melançon, P. (1999). GBF1: A novel Golgi-associated BFA-resistant guanine nucleotide exchange factor that displays specificity for ADP-ribosylation factor 5. J. Cell Biol. 146, 71–84.

Cronin, T.C., DiNitto, J.P., Czech, M.P., and Lambright, D.G. (2004). Structural determinants of phosphoinositide selectivity in splice variants of Grp1 family PH domains. EMBO J. 23, 3711–3720.

Das, S., Malaby, A.W., Nawrotek, A., Zhang, W., Zeghouf, M., Maslen, S., Skehel, M., Chakravarthy, S., Irving, T.C., Bilsel, O., et al. (2019). Structural Organization and Dynamics of Homodimeric Cytohesin Family Arf GTPase Exchange Factors in Solution and on Membranes. Structure 27, 1782–1797.e7.

Dechant, R., Saad, S., Ibáñez, A.J., and Peter, M. (2014). Cytosolic pH regulates cell growth through distinct GTPases, Arf1 and Gtr1, to promote Ras/PKA and TORC1 activity. Mol. Cell 55, 409–421.

Delprato, A., and Lambright, D.G. (2007). Structural basis for Rab GTPase activation by VPS9 domain exchange factors. Nat. Struct. Mol. Biol. 14, 406–412.

DiNitto, J.P., Delprato, A., Gabe Lee, M.-T., Cronin, T.C., Huang, S., Guilherme, A., Czech, M.P., and Lambright, D.G. (2007). Structural basis and mechanism of autoregulation in 3-phosphoinositide-dependent Grp1 family Arf GTPase exchange factors. Mol. Cell 28, 569–583.

Donaldson, J.G., and Jackson, C.L. (2011). ARF family G proteins and their regulators: roles in membrane transport, development and disease. Nat. Rev. Mol. Cell Biol. 12, 362–375.

Drin, G., and Antonny, B. (2010). Amphipathic helices and membrane curvature. FEBS Lett. 584, 1840–1847.

Duronio, R.J., Jackson-Machelski, E., Heuckeroth, R.O., Olins, P.O., Devine, C.S., Yonemoto, W., Slice, L.W., Taylor, S.S., and Gordon, J.I. (1990). Protein N-myristoylation in Escherichia coli: reconstitution of a eukaryotic protein modification in bacteria. Proc. Natl. Acad. Sci. U. S. A. 87, 1506–1510.

Emsley, P., Lohkamp, B., Scott, W.G., and Cowtan, K. (2010). Features and development of Coot. Acta Crystallogr. D Biol. Crystallogr. 66, 486–501.

Franco, M., Chardin, P., Chabre, M., and Paris, S. (1995). Myristoylation of ADP-ribosylation factor 1 facilitates nucleotide exchange at physiological Mg2+ levels. J. Biol. Chem. 270, 1337–1341.

Franzusoff, A., Redding, K., Crosby, J., Fuller, R.S., and Schekman, R. (1991). Localization of components involved in protein transport and processing through the yeast Golgi apparatus. J. Cell Biol. 112, 27–37.

Galindo, A., Soler, N., McLaughlin, S.H., Yu, M., Williams, R.L., and Munro, S. (2016). Structural Insights into Arl1-Mediated Targeting of the Arf-GEF BIG1 to the trans-Golgi. Cell Rep. 16, 839–850.

Gillingham, A.K., and Munro, S. (2007). The small G proteins of the Arf family and their regulators. Annu. Rev. Cell Dev. Biol. 23, 579–611.

Goldberg, J. (1998). Structural basis for activation of ARF GTPase: mechanisms of guanine nucleotide exchange and GTP-myristoyl switching. Cell 95, 237–248.

Grebe, M., Gadea, J., Steinmann, T., Kientz, M., Rahfeld, J.U., Salchert, K., Koncz, C., and Jürgens, G. (2000). A conserved domain of the arabidopsis GNOM protein mediates subunit interaction and cyclophilin 5 binding. Plant Cell 12, 343–356.

Gureasko, J., Galush, W.J., Boykevisch, S., Sondermann, H., Bar-Sagi, D., Groves, J.T., and Kuriyan, J. (2008). Membrane-dependent signal integration by the Ras activator Son of sevenless. Nat. Struct. Mol. Biol. 15, 452–461.

Gustafson, M.A. (2017). The Arf-GEFs Gea1 and Gea2 integrate signals to coordinate vesicle trafficking at the Golgi complex.

Gustafson, M.A., and Fromme, J.C. (2017). Regulation of Arf activation occurs via distinct mechanisms at early and late Golgi compartments. Mol. Biol. Cell 28, 3660–3671.

Halaby, S.L., and Fromme, J.C. (2018). The HUS box is required for allosteric regulation of the Sec7 Arf-GEF. J. Biol. Chem. 293, 6682–6691.

Haun, R.S., Tsai, S.C., Adamik, R., Moss, J., and Vaughan, M. (1993). Effect of myristoylation on GTP-dependent binding of ADP-ribosylation factor to Golgi. J. Biol. Chem. 268, 7064–7068.

Isomura, M., Kikuchi, A., Ohga, N., and Takai, Y. (1991). Regulation of binding of rhoB p20 to membranes by its specific regulatory protein, GDP dissociation inhibitor. Oncogene 6, 119–124.

Jumper, J., Evans, R., Pritzel, A., Green, T., Figurnov, M., Ronneberger, O., Tunyasuvunakool, K., Bates, R., Žídek, A., Potapenko, A., et al. (2021). Highly accurate protein structure prediction with AlphaFold. Nature 596, 583–589.

Kahn, R.A., and Gilman, A.G. (1986). The protein cofactor necessary for ADP-ribosylation of Gs by cholera toxin is itself a GTP binding protein. J. Biol. Chem. 261, 7906–7911.

Kahn, R.A., Goddard, C., and Newkirk, M. (1988). Chemical and immunological characterization of the 21-kDa ADP-ribosylation factor of adenylate cyclase. J. Biol. Chem. 263, 8282–8287.

Kahn, R.A., Randazzo, P., Serafini, T., Weiss, O., Rulka, C., Clark, J., Amherdt, M., Roller, P., Orci, L., and Rothman, J.E. (1992). The amino terminus of ADP-ribosylation factor (ARF) is a critical determinant of ARF activities and is a potent and specific inhibitor of protein transport. J. Biol. Chem. 267, 13039–13046.

Kumari, S., and Mayor, S. (2008). ARF1 is directly involved in dynamin-independent endocytosis. Nat. Cell Biol. 10, 30–41.

Lauer, J., Segeletz, S., Cezanne, A., Guaitoli, G., Raimondi, F., Gentzel, M., Alva, V., Habeck, M., Kalaidzidis, Y., Ueffing, M., et al. (2019). Auto-regulation of Rab5 GEF activity in Rabex5 by allosteric structural changes, catalytic core dynamics and ubiquitin binding. Elife 8.

Lee, J.Y., Chen, H., Liu, A., Alba, B.M., and Lim, A.C. (2017). Auto-induction of Pichia pastoris AOX1 promoter for membrane protein expression. Protein Expr. Purif. 137, 7–12.

Liebschner, D., Afonine, P.V., Baker, M.L., Bunkóczi, G., Chen, V.B., Croll, T.I., Hintze, B., Hung, L.W., Jain, S., McCoy, A.J., et al. (2019). Macromolecular structure determination using X-rays, neutrons and electrons: recent developments in Phenix. Acta Crystallogr D Struct Biol 75, 861–877.

Liu, Y., Kahn, R.A., and Prestegard, J.H. (2010). Dynamic structure of membrane-anchored Arf*GTP. Nat. Struct. Mol. Biol. 17, 876–881.

Lowery, J., Szul, T., Styers, M., Holloway, Z., Oorschot, V., Klumperman, J., and Sztul, E. (2013). The Sec7 guanine nucleotide exchange factor GBF1 regulates membrane recruitment of BIG1 and BIG2 guanine nucleotide exchange factors to the trans-Golgi network (TGN). J. Biol. Chem. 288, 11532–11545.

Malaby, A.W., van den Berg, B., and Lambright, D.G. (2013). Structural basis for membrane recruitment and allosteric activation of cytohesin family Arf GTPase exchange factors. Proc. Natl. Acad. Sci. U. S. A. 110, 14213–14218.

Malaby, A.W., Das, S., Chakravarthy, S., Irving, T.C., Bilsel, O., and Lambright, D.G. (2018). Structural Dynamics Control Allosteric Activation of Cytohesin Family Arf GTPase Exchange Factors. Structure 26, 106–117.e6.

Mastronarde, D.N. (2005). Automated electron microscope tomography using robust prediction of specimen movements. J. Struct. Biol. 152, 36–51.

McDonold, C.M., and Fromme, J.C. (2014). Four GTPases differentially regulate the Sec7 Arf-GEF to direct traffic at the trans-golgi network. Dev. Cell 30, 759–767.

Meissner, J.M., Bhatt, J.M., Lee, E., Styers, M.L., Ivanova, A.A., Kahn, R.A., and Sztul, E. (2018). The ARF guanine nucleotide exchange factor GBF1 is targeted to Golgi membranes through a PIP-binding domain. J. Cell Sci. 131.

Monetta, P., Slavin, I., Romero, N., and Alvarez, C. (2007). Rab1b interacts with GBF1 and modulates both ARF1 dynamics and COPI association. Mol. Biol. Cell 18, 2400–2410.

Mouratou, B., Biou, V., Joubert, A., Cohen, J., Shields, D.J., Geldner, N., Jürgens, G., Melançon, P., and Cherfils, J. (2005). The domain architecture of large guanine nucleotide exchange factors for the small GTP-binding protein Arf. BMC Genomics 6, 20.

Nakane, T., Kimanius, D., Lindahl, E., and Scheres, S.H. (2018). Characterisation of molecular motions in cryo-EM single-particle data by multi-body refinement in RELION. Elife 7.

Novick, P., Ferro, S., and Schekman, R. (1981). Order of events in the yeast secretory pathway. Cell 25, 461–469.

Paczkowski, J.E., and Fromme, J.C. (2016). Analysis of Arf1 GTPase-Dependent Membrane Binding and Remodeling Using the Exomer Secretory Vesicle Cargo Adaptor. Methods Mol. Biol. 1496, 41–53.

Paris, S., Béraud-Dufour, S., Robineau, S., Bigay, J., Antonny, B., Chabre, M., and Chardin, P. (1997). Role of protein-phospholipid interactions in the activation of ARF1 by the guanine nucleotide exchange factor Arno. J. Biol. Chem. 272, 22221–22226.

Park, S.-K., Hartnell, L.M., and Jackson, C.L. (2005). Mutations in a highly conserved region of the Arf1p activator GEA2 block anterograde Golgi transport but not COPI recruitment to membranes. Mol. Biol. Cell 16, 3786–3799.

Peyroche, A., Paris, S., and Jackson, C.L. (1996). Nucleotide exchange on ARF mediated by yeast Gea1 protein. Nature 384, 479–481.

Pocognoni, C.A., Viktorova, E.G., Wright, J., Meissner, J.M., Sager, G., Lee, E., Belov, G.A., and Sztul, E. (2018). Highly conserved motifs within the large Sec7 ARF guanine nucleotide exchange factor GBF1 target it to the Golgi and are critical for GBF1 activity. Am. J. Physiol. Cell Physiol. 314, C675–C689.

Punjani, A., Rubinstein, J.L., Fleet, D.J., and Brubaker, M.A. (2017). cryoSPARC: algorithms for rapid unsupervised cryo-EM structure determination. Nat. Methods 14, 290–296.

Ramaen, O., Joubert, A., Simister, P., Belgareh-Touzé, N., Olivares-Sanchez, M.C., Zeeh, J.-C., Chantalat, S., Golinelli-Cohen, M.-P., Jackson, C.L., Biou, V., et al. (2007). Interactions between conserved domains within homodimers in the BIG1, BIG2, and GBF1 Arf guanine nucleotide exchange factors. J. Biol. Chem. 282, 28834–28842.

Renault, L., Guibert, B., and Cherfils, J. (2003). Structural snapshots of the mechanism and inhibition of a guanine nucleotide exchange factor. Nature 426, 525–530.

Richardson, B.C., and Fromme, J.C. (2012). Autoregulation of Sec7 Arf-GEF activity and localization by positive feedback. Small GTPases 3, 240–243.

Richardson, B.C., and Fromme, J.C. (2015). Biochemical methods for studying kinetic regulation of Arf1 activation by Sec7. Methods Cell Biol. 130, 101–126.

Richardson, B.C., McDonold, C.M., and Fromme, J.C. (2012). The Sec7 Arf-GEF is recruited to the trans-Golgi network by positive feedback. Dev. Cell 22, 799–810.

Richardson, B.C., Halaby, S.L., Gustafson, M.A., and Fromme, J.C. (2016). The Sec7 N-terminal regulatory domains facilitate membrane-proximal activation of the Arf1 GTPase. Elife 5.

Robinson, J.S., Klionsky, D.J., Banta, L.M., and Emr, S.D. (1988). Protein sorting in Saccharomyces cerevisiae: isolation of mutants defective in the delivery and processing of multiple vacuolar hydrolases. Mol. Cell. Biol. 8, 4936–4948.

Soldati, T., Shapiro, A.D., Svejstrup, A.B., and Pfeffer, S.R. (1994). Membrane targeting of the small GTPase Rab9 is accompanied by nucleotide exchange. Nature 369, 76–78.

Sondermann, H., Soisson, S.M., Boykevisch, S., Yang, S.-S., Bar-Sagi, D., and Kuriyan, J. (2004). Structural analysis of autoinhibition in the Ras activator Son of sevenless. Cell 119, 393–405.

Spang, A., Herrmann, J.M., Hamamoto, S., and Schekman, R. (2001). The ADP ribosylation factor-nucleotide exchange factors Gea1p and Gea2p have overlapping, but not redundant functions in retrograde transport from the Golgi to the endoplasmic reticulum. Mol. Biol. Cell 12, 1035–1045.

Su, M.-Y., Fromm, S.A., Zoncu, R., and Hurley, J.H. (2020). Structure of the C9orf72 ARF GAP complex that is haploinsufficient in ALS and FTD. Nature 585, 251–255.

Terwilliger, T.C., Ludtke, S.J., Read, R.J., Adams, P.D., and Afonine, P.V. (2020). Improvement of cryo-EM maps by density modification. Nat. Methods 17, 923–927.

Togawa, A., Morinaga, N., Ogasawara, M., Moss, J., and Vaughan, M. (1999). Purification and cloning of a brefeldin A-inhibited guanine nucleotide-exchange protein for ADP-ribosylation factors. J. Biol. Chem. 274, 12308–12315.

Wang, R., Wang, Z., Wang, K., Zhang, T., and Ding, J. (2016). Structural basis for targeting BIG1 to Golgi apparatus through interaction of its DCB domain with Arl1. J. Mol. Cell Biol. 8, 459–461.

Weiss, O., Holden, J., Rulka, C., and Kahn, R.A. (1989). Nucleotide binding and cofactor activities of purified bovine brain and bacterially expressed ADP-ribosylation factor. J. Biol. Chem. 264, 21066–21072.

Wilfling, F., Thiam, A.R., Olarte, M.-J., Wang, J., Beck, R., Gould, T.J., Allgeyer, E.S., Pincet, F., Bewersdorf, J., Farese, R.V., Jr, et al. (2014). Arf1/COPI machinery acts directly on lipid droplets and enables their connection to the ER for protein targeting. Elife 3, e01607.

Yu, B., Martins, I.R.S., Li, P., Amarasinghe, G.K., Umetani, J., Fernandez-Zapico, M.E., Billadeau, D.D., Machius, M., Tomchick, D.R., and Rosen, M.K. (2010). Structural and energetic mechanisms of cooperative autoinhibition and activation of Vav1. Cell 140, 246–256.

Zhang, Z., Zhang, T., Wang, S., Gong, Z., Tang, C., Chen, J., and Ding, J. (2014). Molecular mechanism for Rabex-5 GEF activation by Rabaptin-5. Elife 3.

Zheng, S.Q., Palovcak, E., Armache, J.-P., Verba, K.A., Cheng, Y., and Agard, D.A. (2017). MotionCor2: anisotropic correction of beam-induced motion for improved cryo-electron microscopy. Nat. Methods 14, 331–332.

Zivanov, J., Nakane, T., Forsberg, B.O., Kimanius, D., Hagen, W.J., Lindahl, E., and Scheres, S.H. (2018). New tools for automated high-resolution cryo-EM structure determination in RELION-3. Elife 7.

Zivanov, J., Nakane, T., and Scheres, S.H.W. (2019). A Bayesian approach to beam-induced motion correction in cryo-EM single-particle analysis. IUCrJ 6, 5–17.

Zivanov, J., Nakane, T., and Scheres, S.H.W. (2020). Estimation of high-order aberrations and anisotropic magnification from cryo-EM data sets in −3.1. IUCrJ 7, 253–267.

